# Microglial Foxo3 shapes dopaminergic vulnerability in Parkinson’s disease

**DOI:** 10.64898/2026.05.22.727098

**Authors:** Texier Baptiste, Roudaud Sarah, Husson Quentin, Prime Morgane, Alma Marie, Dominguez-Coronado Lisaydi, Frieser David, Le Dorze Anne-Louise, Zahm Margot, Iscache Anne-Laure, Alvarez-Fischer Daniel, S. Dejean Anne, Klonjkowski Bernard, Hunot Stéphane, Fruchon Séverine, Szelechowski Marion

**Affiliations:** Univ Toulouse, INSERM, CNRS, Infinity – Toulouse Institute for Infectious and Inflammatory Diseases, Toulouse, France; Institute of Neurogenetics and Institute of Systems Motor Science, University of Lübeck, Lübeck, Germany; Ecole nationale vétérinaire d’Alfort, Anses, INRAE, Laboratoire de Santé Animale, VIROLOGIE, Maisons-Alfort, F-94700, France; Center for Interdisciplinary Research in Biology (CIRB), Collège de France, CNRS UMR 7241, INSERM U1050, Université PSL, Paris, France

## Abstract

Neurodegenerative diseases such as Parkinson’s disease (PD) result from complex interactions between neuronal stress and the surrounding tissue environment, yet the determinants that govern this interplay remain incompletely understood. While neuronal responses to mitochondrial dysfunction and proteotoxic stress have been extensively characterized, the contribution of the microglial state to disease progression remains unclear.

Here, we identify the transcription factor Foxo3 as a key regulator of dopaminergic vulnerability acting predominantly through microglia-, rather than neuron-intrinsic, mechanisms. Foxo3 was rapidly induced and translocated to the nucleus in dopaminergic-like cells in response to mitochondrial complex I inhibition and α-synuclein aggregation, indicating activation of a conserved neuronal stress response. However, neuron-specific deletion of Foxo3 attenuated early Parkinson-like transcriptional signatures but did not confer sustained *in vivo* neuroprotection.

In contrast, microglia-specific deletion of Foxo3 provided robust and durable protection against dopaminergic degeneration across complementary mouse models, including both MPTP intoxication and α-synuclein–driven pathology. Translatomic profiling revealed that Foxo3 deficiency induces extensive transcriptional remodeling of microglia at baseline, establishing a distinct state enriched in immune-related and phagocytic pathways that remains compatible with tissue homeostasis. Strikingly, neuroprotection occurred despite minimal transcriptional reprogramming following neurotoxic insult, suggesting that disease outcome is dictated by the baseline microglial state rather than by the magnitude of transcriptional responses to neurotoxic insult.

Mechanistically, neuroprotection correlated with selective modulation of discrete signaling nodes, including reinforcement of the TREM2-TYROBP axis and attenuation of pathways involved in intracellular signal amplification, rather than with broad suppression of inflammatory programs. These findings indicate that Foxo3 does not primarily regulate the magnitude of microglial activation, but instead defines the baseline configuration and response thresholds that shape subsequent neuroimmune dynamics.

Together, our results identify baseline microglial state as a determinant of neurodegenerative trajectory and position Foxo3 as a central regulator of this process. More broadly, these findings provide a framework in which microglial state-setting, rather than the magnitude of reactive responses alone, governs dopaminergic vulnerability, with potential implications for therapeutic strategies targeting neuroimmune interactions in Parkinson’s disease.

## Introduction

Parkinson’s disease (PD) is a chronic, age-associated neurodegenerative disorder characterized by the progressive loss of dopaminergic neurons (DNs), mainly in the substantia nigra *pars compacta* (SNpc), and the fibrillar aggregation of α-synuclein (aSyn), building up intracytoplasmic inclusions known as Lewy bodies (Holdorff et al., 2013). Beyond these pathological hallmarks, PD is increasingly recognized as a multifactorial disorder in which neuronal vulnerability results from the interplay between intrinsic metabolic stress (neuron-autonomous mechanisms) and non-cell-autonomous mechanisms (Bloem et al., 2021; Kalia & Lang, 2015; Poewe et al., 2017). The disease exhibits substantial etiological heterogeneity, involving a variable contribution of genetic and environmental risk factors. In over 90% of cases, PD occurs sporadically and results from a complex combination of genetic predispositions and environmental exposures, including mitochondrial toxins such as pesticides (Bloem et al., 2021; Nalls et al., 2019). In parallel, monogenic forms of PD have identified key genes, including SNCA, GBA1, PRKN, PINK1, DJ-1 and LRRK2, that converge on pathways regulating mitochondrial homeostasis, proteostasis and cellular stress responses (Blauwendraat et al., 2020; Bonifati et al., 2003; Kitada et al., 1998; Polymeropoulos et al., 1997; Valente et al., 2004). These observations position mitochondrial dysfunction as a central pathogenic hub across both sporadic and familial PD, yet they do not fully explain the selective vulnerability of specific neuronal populations.

A defining feature of SNpc DNs is their autonomous pacemaker activity, which relies in part on sustained Ca^2+^ influx through L-type voltage-gated calcium channels. While Ca^2+^ entry follows its electrochemical gradient, it imposes a substantial energetic burden due to the need for continuous buffering and extrusion, processes that depend on mitochondrial function (Guzman et al., 2010; Surmeier, 2007). In SNpc neurons, unlike those of the ventral tegmental area (VTA), this sustained Ca^2+^ load increases oxidative stress and contributes to selective vulnerability. In addition, their exceptionally large and highly branched axonal arbor imposes considerable energetic demands, requiring a dense mitochondrial network operating near maximal respiratory capacity (Pacelli et al., 2015; Pissadaki & Bolam, 2013). Dopamine metabolism further generates reactive intermediates that impair mitochondrial function (Burbulla et al., 2017; Surmeier, 2018), while pathological aSyn assemblies directly disrupt mitochondrial integrity (Grünewald et al., 2019). Together, these features place SNpc neurons at the edge of metabolic homeostasis and particularly sensitive to mitochondrial perturbations.

Beyond neuron-intrinsic mechanisms, PD pathophysiology critically involves non-cell-autonomous processes, particularly neuroinflammation. Injured neurons release immunogenic signals, including misfolded aSyn species, that activate microglia and trigger immune responses (Heneka et al., 2015; Ransohoff, 2016; Tansey & Romero-Ramos, 2019). Microglial activation is detectable at early disease stages and is characterized by proliferation, phenotypic remodeling and increased inflammatory signaling (Smajić et al., 2022). Activated microglia can adopt neurotoxic phenotypes, releasing cytokines and reactive oxygen species that further damage neurons and promote astrocyte reactivity (Hong et al., 2016; Liddelow et al., 2017; Szepesi et al., 2018). In parallel, increasing evidence supports a role for adaptive immunity in Parkinson’s disease. T cell infiltration has been consistently observed in the PD brain, including in early disease stages (Galiano-Landeira et al., 2020), while antigen-specific T cell responses directed against aSyn and other neuronal antigens further support the existence of antigen-driven mechanisms (Lindestam Arlehamn et al., 2020; Sulzer et al., 2017; Williams et al., 2024). Collectively, these findings highlight a dynamic interplay between innate and adaptive immune responses in PD, yet how these immune components integrate with neuronal stress pathways to shape disease progression remains unresolved. Emerging evidence further suggests that microglial responses are not solely determined by the nature of the insult, but are critically shaped by their pre-existing functional configuration, which can durably shape subsequent responses to injury through mechanisms akin to innate immune memory (Netea et al., 2020; Wendeln et al., 2018).

Recent work further indicates that microglial metabolic state is a key determinant of their functional phenotype and inflammatory output. In particular, mitochondrial function governs microglial activation states and their interaction with neurons, positioning immunometabolism as a central regulator of neuroinflammation (Baik et al., 2019; Lauro & Limatola, 2020). These observations raise the possibility that shared metabolic regulators coordinate neuronal vulnerability and immune responses within the diseased brain.

Among transcriptional regulators integrating metabolic and stress signals, the Forkhead box O (Foxo) family, and in particular Foxo3, has emerged as a critical node linking mitochondrial function, redox homeostasis and immune regulation (Webb & Brunet, 2014). Foxo3 is activated in response to oxidative stress, nutrient deprivation and growth factor withdrawal, and regulates key cellular processes including autophagy, metabolism and cell survival (Brunet et al., 1999; Webb & Brunet, 2014). Several lines of evidence suggest a functional connection between Foxo3 and PD-related pathways. Foxo3 regulates mitophagy and intersects with pathways controlling mitochondrial quality, including the PINK1/Parkin axis (Pickrell & Youle, 2015; Ryan et al., 2015). Importantly, its activity has been linked to DN fate and aSyn accumulation, including in one of the few mammalian PD models directly addressing Foxo3 function (Pino et al., 2014). In addition, allele-specific FOXO3 binding at the SNCA locus in human DNs models supports its involvement in neuron-autonomous pathogenic mechanisms (Chang et al., 2017; Prahl et al., 2023).

In parallel to its neuron-autonomous functions, Foxo3 is increasingly recognized as an immunometabolic regulator. Foxo3 controls inflammatory mediator production in myeloid cells and regulates the polarization and persistence of CD4^+^ T lymphocytes (Dejean et al., 2009; Joulia et al., 2024; Stienne et al., 2016). In the CNS, Foxo3 is a central regulator of cellular stress responses and immune-related transcriptional programs. However, its specific role in microglial biology and neuroinflammation remains poorly defined. Early studies based on *in vitro* systems suggested a potential involvement in oxidative stress responses and inflammatory signaling (Shang et al., 2009b, 2009a; Webb & Brunet, 2014), but *in vivo* evidence remains limited. More recently, transcriptomic analyses in human disease have identified Foxo3 as a candidate regulator of microglial state transitions, including inflammatory and interferon-associated programs (Sun et al., 2023). These findings extend previous work defining disease-associated microglial states in neurodegeneration (Deczkowska et al., 2018; Keren-Shaul et al., 2017), and suggest that Foxo3 may contribute to the transcriptional regulation of these programs.

Together, these observations position Foxo3 at the interface between neuronal stress and neuroimmune regulation. However, whether Foxo3 primarily promotes neuronal vulnerability, supports adaptive responses, or shapes neuroimmune crosstalk in PD remains unclear. In particular, whether pre-existing microglial states actively determine the trajectory of neurodegeneration, rather than merely reflecting neuronal injury, remains unresolved. Here, we addressed this question by combining constitutive and cell type-specific genetic deletion strategies with complementary toxin- and aSyn-based models of PD.

## Materials et Methods

### In vitro experiments

#### Cell culture and maintenance of SH-SY5Y cells

Human neuroblastoma SH-SY5Y cells (Sigma-Aldrich) were maintained in Dulbecco’s Modified Eagle Medium (DMEM) containing 4.5 g/L glucose, supplemented with 10% fetal bovine serum, 8 mM L-glutamine, 1 mM sodium pyruvate, 50 U/mL penicillin, and 50 µg/mL streptomycin. Cells were cultured at 37 °C in a humidified incubator with 5% CO_2_. Cells were passaged twice weekly using trypsin/EDTA, and experiments were performed using cells at passage ≤ 15 to minimize drift. For experimental treatments and subsequent immunocytochemistry, cells were seeded one day prior to treatment on glass coverslips in 24-well plates (7.5 × 10^4^ cells/well in 500 µL medium), reaching 60-70% confluence at the start of treatment.

#### MPP^+^ treatments

MPP^+^ iodide (Sigma-Aldrich) was resuspended as a 33 mM stock solution in ultrapure water, aliquoted, and stored at −20 °C. Aliquots were thawed only once and used within 4 h. For treatments, MPP^+^ was diluted immediately before use in serum-free, pyruvate-free DMEM pre-warmed to 37 °C.

#### CAV2-hSynTR110/EGFP transduction

CAV2 vectors were E1-deleted (ΔE1) and are replication-deficient (Klonjkowski et al., 1997). Vector production, titration, and quality control were performed by the laboratory of Dr. Bernard Klonjkowski (École Nationale Vétérinaire d’Alfort, Maisons-Alfort, France). Viral genome copy number was determined by quantitative PCR, and infectious titers (50% tissue culture infectious dose, TCID_50_) were calculated using the Reed-Muench method.

For viral transduction experiments, SH-SY5Y cells were seeded as described above and incubated with CAV2-hSyn^TR110^/EGFP vectors diluted in complete culture medium. Viral aliquots were thawed once and used immediately. The viral dose was empirically determined to achieve approximately 75% transduction efficiency after 48 h of incubation, from a stock concentration of 1.25 x 10^5^ TCID_50_ /µL.

#### Immunofluorescence staining

For immunocytochemistry, cells grown on glass coverslips were fixed at the end of the experiment with 4% PFA, washed with DPBS and stored at 4 °C in DPBS containing 0.1% sodium azide for up to one week before staining.

Cells were permeabilized with 0.25% Triton X-100 in DPBS for 15 min, followed by blocking in DPBS containing 0.25% Triton X-100 and 3% bovine serum albumin (BSA) for 1 h at room temperature. Coverslips were incubated overnight at 4 °C with primary antibodies diluted in blocking buffer: anti-cleaved caspase-3 (1:500, Millipore), anti-Foxo3 (1:300, Invitrogen), anti-α-synuclein (syn204, 1:100, Cell Signaling). After extensive washing, cells were incubated for 1 h at room temperature with Alexa Fluor-conjugated secondary antibodies (1:500, Invitrogen). Nuclei were counterstained with DAPI (1:50,000, BD Biosciences) for 5 min. Coverslips were washed and mounted using Mowiol 4-88 mounting medium (Sigma-Aldrich).

#### Confocal microscopy and image analysis

Fluorescent images were acquired using a Leica SP8 confocal microscope equipped with a 20× objective. For each coverslip, four fields of view were selected in a blinded manner using a predefined acquisition grid to avoid sampling bias. For each field, z-stacks consisting of 16 optical sections (0.68 µm step size) were acquired to capture the full cellular volume.

Acquisition parameters (laser power, detector gain, pinhole, and resolution) were optimized to prevent signal saturation and were strictly maintained constant across all conditions within each experiment. All samples from a given experiment were imaged during the same acquisition session to minimize inter-run variability. Image analysis was performed using ImageJ (Fiji). Maximum intensity projections were generated for all z-stacks prior to analysis. Background subtraction was performed using identical parameters across all images. Cell segmentation was performed based on DAPI staining to define nuclear regions. Cytoplasmic regions were defined by subtracting the nuclear mask from the total cell area, when appropriate. Cells with overlapping nuclei or segmentation artifacts were excluded from analysis. Pyknotic nuclei were identified based on increased DAPI intensity and reduced nuclear area, using consistent thresholding parameters across all conditions. Mean fluorescence intensity was quantified at the single-cell level. Cleaved caspase-3 and Foxo3 signals were measured per cell and normalized to the mean intensity of control conditions when indicated. Nuclear and cytoplasmic fluorescence intensities were distinguished using DAPI-based nuclear masks, allowing calculation of nuclear-to-cytoplasmic ratios.

For each experimental condition, 200-500 cells were analyzed per independent experiment. The mean value obtained per experiment was used as a biological replicate for statistical analysis.

### Animal models

#### Mice

All procedures described herein were conducted in accordance with European ethical regulations, approved by the local Animal Experimentation Ethics Committee (CEEA-122), and authorized by the French Ministry of Higher Education, Research and Innovation (project authorization number APAFIS#23457-2019122710273327v3). Animals used in this study were C57BL/6 mice maintained under pathogen-free conditions in the breeding facility of CREFRE (Toulouse UMS06). Animals were housed under constant temperature (22±1 °C), relative humidity (55±1%), and 12 h light/12 h dark cycle (lights on at 7:00 am). Mice had ad libitum access to tap water and commercial rodent chow.

Foxo3-deficient C57BL/6 mice were generated using embryonic stem (ES) cell clones from the OmniBank® ES cell library of randomly targeted cell lines (Lexicon Genetics; used as previously described (Dejean et al., 2009)). This line was maintained by breeding heterozygous Foxo3^+/-^ reproducers, and Foxo3^+/+^ littermates were used as wild-type (WT) controls.

To generate lines with dopaminergic-neuron-specific (Foxo3^ΔDN^) or microglia-specific (Foxo3^ΔmG^) deletion of Foxo3, a Foxo3^fl/fl^ C57BL/6 line (Jackson #037528; (Castrillon et al., 2003)) was used, and crossbred with *Dat*^*iCre*^ (Turiault et al., 2007) (kind gift from Dr. Tricoire, Neuroscience Paris Seine) and *Cx3cr1*^*CreERT2*^ (Cx3cr1tm2.1(cre/ERT2)Litt; MGI:5617710; used as previously described (Yona et al., 2013)) lines respectively. Those lines were further crossed with a Rpl22^Haflox/+^ (RiboTag) line (Jackson Laboratory, #029977; (Sanz et al., 2009)) in order to induce expression of ribosomal HA-tag in dopaminergic neurons and microglia for subsequent specific RNA sequencing.

For dopaminergic neuron-specific deletion, Foxo3^fl/fl^ mice were crossed with *Dat*^*iCre*^ mice, in which Cre recombinase is expressed under the control of the dopamine transporter (Slc6a3) promoter. This results in constitutive deletion of Foxo3 in dopaminergic neurons (Foxo3^ΔDN^) without the need for temporal induction.

For inducible deletion in resident myeloid cells, Foxo3^fl/fl^ mice were crossed with *Cx3cr1*^*CreERT2*^ mice, in which Cre recombinase is expressed under the control of the CX3CR1 promoter and fused to a tamoxifen-inducible estrogen receptor (ERT2), allowing temporal control of recombination (Foxo3^ΔmG^).

Only male animals were examined in the present study to avoid ambiguity associated with reported sex-related differences.

#### Tamoxifen-induced recombination and validation

Tamoxifen was dissolved in Miglyol and administered intraperitoneally at a dose of 2 mg per injection (200 µL per mouse) once daily for five consecutive days to induce recombination in CX3CR1-expressing cells. Tamoxifen injections were performed in adult mice aged 6–8 weeks.

Following tamoxifen administration, a delay period was implemented to allow the turnover of short-lived circulating CX3CR1^+^ peripheral myeloid cells, while long-lived resident microglia retained recombination. All Parkinson’s disease models were induced at least 28 days after the last tamoxifen injection.

Recombination efficiency was assessed 21 days after tamoxifen administration by flow cytometry on brain and peripheral lymphoid tissues. Foxo3 expression was quantified at the protein level by intracellular staining and was reduced by more than 80% in Foxo3^ΔmG^ mice compared to Foxo3^WT^ controls. In contrast, Foxo3 expression levels in peripheral myeloid cells from cervical lymph nodes were comparable between genotypes at baseline, indicating that recombination in short-lived circulating CX3CR1^+^ populations was transient and had largely resolved by this time point (Additional file 10: Fig. S10).

These results confirm that, at the time of PD induction, Foxo3 deletion is efficiently restricted to resident CNS myeloid cells.

#### MPTP mouse model

9-week-old male mice were housed in A2-level animal facility and handled daily for a period of 10 days prior to protocol induction for habituation. Animals weighed between 24 and 29 g before the protocol. Less than 24 h before injection, MPTP-HCl powder (Sigma) was resuspended in DPBS buffer and kept cool and away from light before use. MPTP was administered intraperitoneally at a dose of 25 mg/kg (free base equivalent, corresponding to 30 mg/kg MPTP-HCl) in a volume of 100 µL per 10 g of body weight. Animals were placed on a heating pad for 2 h following injection for recovery. Injections were repeated every 24 h for 5 consecutive days. Animals were provided with food and water gel and monitored until recovery. Animals were randomly assigned to experimental groups, and all analyses were performed in a blinded manner.

#### HPLC analysis of striatal neurotransmitters and MPP^+^

Mice were deeply anesthetized using a mixture of ketamine and xylazine and striatal tissues were quickly dissected, weighed, and stored at −80 °C until analysis. Samples were thawed on ice and homogenized in ice-cold 0.1 N HClO_4_ (250 µL per sample) using a hand sonicator (3 × 10 s pulses). Homogenates were incubated on ice for 30 min and centrifuged at 13,000 rpm for 20 min at 4 °C. Supernatants were collected, filtered through SpinX tubes, and stored at −80 °C until analysis.

For neurotransmitter quantification, 15 µL of filtrate was mixed with 15 µL of 0.4 N HClO_4_. Dopamine (DA) and DOPAC were separated on a Nucleosil C18 column (125 × 3 mm; pre-column 5 × 4 mm, Knauer) using TEST mobile phase supplemented with 10% (v/v) acetonitrile (flow rate 0.50 mL/min) and quantified by electrochemical detection (+650 mV, 2 nA/V; Ultimate 3000 system). External calibration curves were generated using seven standard concentrations (10^−7^–10^−4^ mol/L; detection limit 10^−9^ mol/L). Neurotransmitter levels were normalized to tissue weight and expressed as nmol per mg of wet tissue.

MPP^+^ levels were measured at 60, 120, and 240 min following intraperitoneal injection of 25 mg/kg MPTP. Tissue processing was performed as described above. MPP^+^ was separated on a Nucleosil 100 C18 column (250 × 4 mm; pre-column 5 × 4 mm, Knauer) and quantified by UV detection (295 nm; Ultimate 3000 system; mobile phase: 2.7 g KH_2_SO_4_ in 697 mL acetonitrile/L, pH 2.5; injection volume: 10 µL).

Recovery and extraction efficiency were monitored using external standards processed in parallel and were consistent across experimental groups.

#### AAV-induced synucleinopathy model

AAV2/7-derived viral vectors were produced and provided by the Leuven Viral Vector Core (KU Leuven, Belgium). Vector constructs included the hybrid CMV-enhanced synapsin promoter to ensure neuronal specificity and encoded either hSyn^TR110^ (the first 110 amino acids of human α-synuclein) or EGFP (control vector).

Nine-week-old male mice (24–29 g) were anesthetized with a ketamine/xylazine mixture and placed in a stereotaxic frame on a heating pad. Local anesthesia (lidocaine) was applied to the ear canals and subcutaneously at the incision site. The skull was aligned horizontally prior to injection. Stereotaxic injections were performed unilaterally into the substantia nigra pars compacta (SNpc) using the following coordinates relative to bregma: AP = −3.1 mm, ML = ±1.2 mm, DV = −4.0 mm. A total volume of 2 µL of viral vector (8 × 10^8^ genome copies (GC)/µL) was injected at a rate of 200 nL/min using a 10 µL Hamilton syringe (Dutscher), corresponding to a total dose of 1.6 × 10^9^ GC per injection site. The needle was left in place for an additional 10 minutes before slow withdrawal to prevent reflux.

Animals received postoperative care including food and hydrogel supplementation and were monitored until full recovery. Mice were sacrificed 8 weeks after injection. Animals were randomly assigned to experimental groups, and all analyses were performed in a blinded manner.

### Brain slice immunostaining

Mice were deeply anesthetized using a mixture of ketamine and xylazine and blood was removed by intra-cardiac perfusion of a saline buffer (DPBS). Saline perfusion was immediately followed by 4% paraformaldehyde (PFA) perfusion. Brains were dissected and post-fixed 24 h in 4% PFA (4 °C), after what they were dehydrated in a hypertonic solution (30% sucrose, 0.2% sodium azide in DPBS, 48 h, 4 °C). Brains were then sectioned using a Leica SM2010 R Microtome to obtain serial 20 µm-thick coronal sections spaced 200 µm apart throughout the regions encompassing the substantia nigra pars compacta and the striatum.

#### TH immunostaining and quantification

Free-floating brain sections were washed in PBS. Endogenous peroxidase activity was quenched by incubating the sections for 5 min at room temperature in a PBS solution containing 20% methanol and 10% hydrogen peroxide solution. Sections were then permeabilized for 5 min at room temperature in PBS containing 0.25% Triton X-100. Non-specific binding sites were blocked by incubating the sections for 30 min at room temperature in a blocking buffer consisting of 0.25% Triton X-100 and 4% bovine serum albumin (BSA) in PBS. Sections were incubated overnight at 4 °C with anti-TH primary antibodies (US Biological, Cat. No. T9237-13; 1:1000) diluted in blocking buffer.

After primary antibody incubation, sections were rinsed in PBS and incubated for 30 min at room temperature with biotinylated goat anti-rabbit secondary antibodies (Invitrogen, Cat. No. 31820; 1:500) prepared in PBS. Sections were then rinsed again and incubated for 1 h at room temperature with avidin–biotin–peroxidase complex (Eurobio Scientifc, VECTASTAIN® Elite ABC-HRP Kit, Peroxidase (Standard), Cat. No. PK-6100; 1:125). Sections were washed in PBS and staining was revealed by incubating the sections for 1 min 45 s under vigorous agitation in 3,3⍰-diaminobenzidine (DAB) solution prepared according to the manufacturer’s instructions (Eurobio Scientifc, ImmPact DAB substrate, Cat. No. SK-4105), after which reaction was stopped immediately by rinsing in PBS. Finally, sections were mounted onto glass slides, dehydrated with ethanol and xylene, and coverslipped using resin mounting medium.

Brightfield images of DAB-TH-immunostained sections were analyzed using QuPath software (v0.4.4). Regions of interest (ROIs) corresponding to the dorsal striatum (DS) were manually delineated on each section based on anatomical landmarks defined in the Allen Mouse Brain Atlas. For each ROI, DAB optical density (OD) was quantified using QuPath’s built-in measurement tools. To correct for non-specific background staining, an additional ROI was defined in the cerebral cortex, a region largely devoid of TH-positive fibers. The OD value measured in the cortical ROI was subtracted from the OD value measured in the ipsilateral dorsal striatum, yielding a background-corrected measure of TH immunoreactivity. For each animal, measurements were performed at six rostrocaudal levels of the dorsal striatum. These levels were selected from 20 µm-thick coronal sections, each separated by 200 µm. The mean of the six background-corrected OD values was calculated and used as an estimate of dopaminergic fiber density in the dorsal striatum.

The mouse SNpc spans approximately 1,500 µm along the rostrocaudal axis (Nelson et al., 1996). Accordingly, seven representative rostrocaudal levels were analyzed from 20-µm-thick coronal sections spaced 200 µm apart, allowing a fair extrapolation of the whole SNpc region. To exclude adjacent TH-positive populations (A8 and A10), the SNpc (A9) was anatomically delineated at each level using QuPath (v0.4.4), based on established anatomical criteria and the Allen Mouse Brain Atlas. Within each SNpc ROI, TH-positive neurons were counted manually using the “Add Points” tool in QuPath. Neurons were included in the analysis only when the cell body was clearly identifiable and fully visible within the section.

#### Neuroinflammation analysis and quantification on brain slices

Free-floating sections were washed sequentially under gentle agitation in PBS and PBS-T (PBS 1X containing 0.25% Triton X-100). Sections were then incubated for 45 min in blocking buffer composed of PBS-T supplemented with 4% bovine serum albumin (BSA) and 3% fetal bovine serum (FBS). Sections were incubated overnight at 4°C under agitation with primary antibodies: anti-GFAP (Dako, Cat. No. Z0334, 1:500), anti-IBA1 (Synaptic Systems, Cat. No. 234009; 1:1,000), anti-CD3 (Abcam, Cat. No. A311089; 1:500), anti-Tyrosine hydroxylase (TH, ImmunoStar, Cat. No. 22941; 1:1,000). The following day, sections were incubated with secondary antibodies coupled with fluorochromes (Invitrogen) for 2 h at room temperature protected from light. Sections were then counterstained with DAPI (BD Biosciences, Cat. No. 564907) for 5 min, washed again in PBS, and mounted onto microscope slides.

Imaging was performed using the Leica TCS SP8 Confocal Microscope at 20X magnification. For each sample, 4 images were acquired in each region of interest (DS or SNpc), and cell quantification was performed using ImageJ (*Analyze particles* function on thresholded images).

### Flow cytometry

#### Sample preparation for flow cytometry

Mice were deeply anesthetized with ketamine/xylazine and transcardially perfused with 20 mL of cold PBS at a constant flow rate (5 mL/min). Brains were rapidly collected and either processed whole or microdissected to isolate the striatum and dorsal mesencephalon (containing the SNpc). Cervical lymph nodes (CLNs) were then collected and mechanically dissociated using a Potter homogenizer.

Tissues were enzymatically dissociated in DMEM containing collagenase D (1 mg/mL; Roche) and DNase I (10 µg/mL; Sigma) for 20 min at 37 °C, with gentle inversions every 5 min. Samples were then mechanically triturated (8-10 passes with a Pasteur pipette), diluted with 5 mL of MACS buffer (PBS supplemented with 0.5% bovine serum albumin (BSA) and 2 mM EDTA) supplemented with 1 mL fetal bovine serum (FBS), and filtered through a 70 µm cell strainer. Cells were washed in MACS buffer and centrifuged at 300 × g for 10 min at 4 °C. Myelin was removed using a 30% isotonic Percoll gradient followed by centrifugation at 800 × g for 20 min at 4 °C without brake. After removal of the myelin layer, cells were washed in PBS, centrifuged at 300 × g for 10 min, and resuspended in MACS buffer for subsequent flow cytometry staining.

#### High-dimensional flow cytometry analysis of microglial states

Single-cell suspensions prepared from brain tissue were first incubated with 1% mouse serum and anti-CD16/32 Fc receptor blocking antibody to reduce non-specific antibody binding. Cells were subsequently stained with LIVE/DEAD™ Fixable Near-IR viability dye followed by fluorochrome-conjugated antibodies directed against surface markers associated with homeostatic microglial identity, phagocytic activity, antigen presentation and immune activation.

The panel included antibodies against CD45, CD11b, CX3CR1, TMEM119, Ly6C, TREM2, CD86, CD80, CD14, MHC-II, TIM-3, CCR2, CD115, CD206, CD38, Clec7A, MerTK, TIM-4, PD-L1, CD64 and CD11c. Detailed information for all antibodies, fluorochromes, clones is provided in Supplementary Table 3.

Data were acquired on a Symphony A5 flow cytometer (Becton Dickinson) and analyzed using OMIQ (Dotmatics) and FlowJo softwares. After exclusion of debris, doublets and dead cells, resident brain myeloid cells were identified based on intermediate-to-high CD45 expression, CD11b, CX3CR1 and TMEM119 expression together with Ly6C exclusion. High-dimensional analyses were performed using UMAP dimensional reduction on concatenated microglial populations from all experimental groups. Marker expression was visualized using feature plots and quantified as percentage of positive cells and/or median fluorescence intensity (MFI), as indicated in the corresponding figure legends. Gating strategies and dimensional reduction parameters were kept identical across experimental groups. Fluorescence minus one (FMO) controls or appropriate negative controls were used when applicable.

### Translatomic analyses

#### Translatome sample preparation

Cell type-specific ribosome-associated mRNAs were isolated using the RiboTag strategy as previously described (Sanz et al., 2009) with minor adaptations. Homogenization buffer consisted of 50 mM Tris-HCl (pH 7.4), 100 mM KCl, 12 mM MgCl_2_, 1% NP-40, supplemented with 1 mM DTT, 100 µg/mL cycloheximide, 100 µg/ml heparin and RNase inhibitors (RNAsin, Promega). High-salt buffer was identical except for KCl adjusted to 300 mM. Specific brain areas from Rpl22^HA^-expressing mice were rapidly dissected without perfusion and homogenized at 2-3% (w/v) in ice-cold homogenization buffer. Lysates were clarified by centrifugation at 10,000 × g for 10 min at 4 °C, and supernatants were collected. An aliquot (50-100 µL) was retained as input and stored at −80 °C.

For immunoprecipitation, 800 µL of clarified lysate were incubated with 2 µL of mouse monoclonal anti-HA antibody (HA.11, Covance) for 4 h at 4 °C under gentle rotation. Dynabeads Protein G magnetic beads (Invitrogen) were washed and added to the antibody– lysate mixture, followed by overnight incubation at 4 °C with rotation. Beads were washed three times with high-salt buffer (50 mM Tris-HCl (pH 7.4), 300 mM KCl, 12 mM MgCl_2_, 1% NP-40, 1 mM DTT, and 100 µg/mL cycloheximide) to reduce non-specific binding.

Ribosome-associated mRNAs were released by adding 350 µL of RLT lysis buffer supplemented with β-mercaptoethanol directly to the beads and vortexing for 30 s. RNA was purified using the RNeasy Plus Micro Kit (Qiagen) according to the manufacturer’s instructions and eluted in 20 µL RNase-free water. RNA concentration and purity (A260/280 and A260/230 ratios) were assessed using a NanoDrop spectrophotometer (Thermo Fisher Scientific). Purified RNA samples were stored at −80 °C until downstream analyses.

All procedures were performed under RNase-free conditions and at 4 °C whenever possible to preserve RNA integrity.

#### Reverse Transcription and Quantitative Real-Time PCR (qPCR)

Prior to reverse transcription, all RNA samples were adjusted to the lowest concentration among the samples using PCR-grade water to ensure uniform input. Complementary DNA (cDNA) synthesis was performed using the SuperScript III First-Strand Synthesis System (Invitrogen), following the manufacturer’s protocol. cDNA concentration and purity was measured using the NanoDrop 1000 spectrophotometer.

Quantitative PCR reactions were performed in 384-well plates (Roche, Cat. No. 04729749001). Each well contained 8 µL of reaction mix composed of 2 µL of 10 µM forward and reverse primers, 1 µL of PCR-grade water, and 5 µL of LightCycler 480 SYBR Green I Master (Roche, Cat. No. 04887352001). Subsequently, 2 µL of cDNA template were added to each well, resulting in a total reaction volume of 10 µL. Primers used for quantitative PCR were as follows: *Foxo3* (forward: 5⍰-CAAACGGCTCACTTTGTCCC-3⍰; reverse: 5⍰-GTTGTGCCGGATGGAGTTCT-3⍰), *Foxo1* (forward: 5⍰-AGCTGGGTGTCAGGCTAAGA-3⍰; reverse: 5⍰-TAAGGAGGGGTGAAGGGCA-3⍰), *Foxo6* (forward: 5⍰-AACCCTCCTCACTGTCTCGG-3⍰; reverse: 5⍰-CGCACAAAGTACTCCAGGTG-3⍰), *Slc6a3* (DAT) (forward: 5⍰-CCACAGATGGACCTGGGTTG-3⍰; reverse: 5⍰-CATGGCACTGTCGATACCCA-3⍰), *Th* (forward: 5⍰-AACCCTCCTCACTGTCTCGG-3⍰; reverse: 5⍰-CGCACAAAGTACTCCAGGTG-3⍰), *Itgam* (forward: 5⍰-CAAATAGCCAGCCTCAGTGC-3⍰; reverse: 5⍰-GAGCCCAGGGGAGAAGTG-3⍰), *Cx3cr1* (forward: 5⍰-AAGTTCCCTTCCCATCTGCT-3⍰; reverse: 5⍰-GGACAGGAAGATGGTTCCAA-3⍰), and *Gapdh* (forward: 5⍰-TGAAGCAGGCATCTGAGG-3⍰; reverse: 5⍰-CGAAGGTGGAAGAGTGGGAG-3⍰).

Amplification was carried out using the LightCycler 480 II (Roche) under the following cycling conditions : initial denaturation at 95 °C for 5 minutes; 40 cycles of 95 °C for 15 seconds, 60 °C for 10 seconds, and 72 °C for 15 seconds; followed by a final step at 40 °C for 30 seconds.

Cycle threshold (Ct) values were normalized to *Gapdh* as the endogenous control. Relative gene expression levels of *Foxo3* were determined by comparing normalized Ct values between experimental conditions.

#### Library preparation and sequencing

Libraries were prepared from 8 ng of RNA using the Illumina® Stranded Total RNA Prep, Ligation with Ribo-Zero Plus kit. Libraries were quantified using a Qubit spectrophotometer and assessment of size distribution and integrity was performed with the TapeStation 4150 (Agilent). Samples were indexed and sequenced (paired-end reads of 150 bp) on the NovaSeq 6000 system (Illumina) from the Genomic and Transcriptomic Platform of the GenoToul (Toulouse).

#### Quantification of RNA expression

Raw sequencing reads were processed using the nf-core RNAseq pipeline v.3.14.0 Nextflow v.24.04.2 (Di Tommaso et al., 2017; Ewels et al., 2020). Briefly, this pipeline trims adapters and removes low quality sequences using Cutadapt v.3.4 wrapped with Trimgalore v.0.6.7 (Krueger, F. (2023): https://www.bioinformatics.babraham.ac.uk/projects/trim_galore/). rRNAs are filtered out using SortMeRNA v.4.3.4 (Kopylova et al., 2012). Then, STAR v.2.7.9a (Dobin et al., 2013) aligns trimmed reads to the Mus musculus GRCm39 assembly. Parameters used to run STAR were:

*--outSAMstrandField intronMotif --quantTranscriptomeBan Singleend --runRNGseed 0*

*--twopassMode Basic --quantMode TranscriptomeSAM*.

Finally, Salmon v.1.10.1 (Patro et al., 2017) quantifies reads with parameter --libType=ISR based on the Mus musculus annotation from Ensembl release 107 (Cunningham et al., 2022).

#### Differential Expression Analyses

Raw Count Matrix was imported on R using the R package DESeq2 v.1.42.1 (Love et al., 2014). Design parameter from DESeqDataSetFromTximport() function was set to ~ condition were condition indicates the genotype and treatment. Expressed genes were identified as genes with at least a sum of 10 raw counts. We used DESeq() and result() functions, with default parameters, to normalize data and run DEA respectively.

PCA were performed using the PCA() function on the top 500 most variable genes after variance stabilizing transformation using the vst() function. Heatmaps were generated using the pheatmap() function from the pheatmap package. Depending on the analysis, heatmaps represented either log2 fold-change values relative to the indicated control condition or row-scaled variance-stabilized normalized expression values (Z-scores). Heatmaps were further adapted using Excel.

#### Enrichment analyses

We ran the enrichment analyses with the R package clusterProfiler v.4.10.1 (Wu et al., 2021) using gene sets from Gene Ontology (GO.db_3.18.0) and KEGG (online version of April 2025) databases. Over Representation Analyses were performed on differentially expressed genes (DEG) defined as genes with an adjusted p-value below 0.05 and an absolute log2 Fold Change greater than 1.

Gene Set Enrichment Analyses were performed on gene list ranked by log2FoldChange extracted from DESeq2 object, using WEB-based GEne SeT AnaLysis Toolkit https://www.webgestalt.org/). Plots were generated using the ClusterProfiler functions dotplot(), treeplot() and gseaplot().

### Public single-cell RNA-sequencing data analysis

Publicly available single-cell RNA-sequencing datasets, including raw count matrices, cluster annotations and dimensional reduction coordinates, were imported and analyzed using the Seurat R package (v5.4.0) (Hao et al., 2021). Data visualization and exploratory analyses were performed using the DimPlot(), DotPlot() and DoHeatmap() functions. Cell population proportion graphs were generated using the ggplot2 R package.

### Statistical analysis

Data are presented as mean ± SEM. Statistical analyses were performed using GraphPad Prism (v10.3.1, GraphPad Software, San Diego, CA, USA). Data are presented as individual values with bars indicating mean ± SEM, unless otherwise stated in the figure legends. Each dot represents one independent biological replicate (one mouse or one independent experiment), except for cell-based imaging quantifications, where each dot represents the mean value obtained from 200-500 cells within one independent experiment.

No statistical method was used to predetermine sample size. Sample sizes were chosen based on prior experience with the corresponding models and were consistent with those commonly used in the field. Animals or samples were not excluded from the analyses unless predefined technical exclusion criteria were met. Investigators were blinded to genotype and treatment during image acquisition and quantification whenever applicable.

For comparisons of experimental groups organized according to two independent variables, typically genotype and treatment, data were analyzed using two-way ANOVA followed by Sidak’s or Tukey’s multiple-comparisons test, as specified in the figure legends. This approach was used for dopaminergic neuron counts, striatal TH optical density, and inflammatory cell quantifications when four groups were compared in parallel (e.g. WT-PBS, WT-MPTP, KO/Δ-PBS, KO/Δ-MPTP).

For comparisons between two independent groups, an unpaired two-tailed Mann–Whitney test was used when only one comparison was performed or when sample size was limited and distribution could not be assumed to be normal. This applied, for example, to comparisons of normalized lesion values between WT and KO mice in the unilateral AAV-hSyn paradigm.

For comparisons to a theoretical reference value (typically 100% for normalized expression values or normalized signal intensity), a two-tailed one-sample Wilcoxon signed-rank test was used. This approach was applied to qRT-PCR validations and to cell-based quantifications normalized to a reference condition.

For analyses in which several experimental conditions were normalized to the mean of a reference group within each independent experiment and then tested against that normalized reference value, a two-tailed one-sample Wilcoxon signed-rank test was used. This was applied to selected neuroinflammatory readouts expressed relative to PBS controls.

For transcriptomic analyses, differential expression was performed on raw count data using the dedicated RNA-sequencing analysis pipeline described in the Methods. Genes were considered significantly differentially expressed according to the thresholds indicated in the corresponding Results section and figure legends (typically adjusted p value < 0.05, with fold-change cutoffs when specified). For Gene Set Enrichment Analysis (GSEA), significance was assessed using normalized enrichment score (NES) and false discovery rate (FDR), as indicated in the figures. For Gene Ontology enrichment analyses, adjusted p values or FDR values were used as provided by the enrichment pipeline. All transcriptomic analyses were performed with investigators blinded to genotype and treatment during data processing.

All tests were two-sided. Exact statistical tests used for each panel are indicated in the corresponding figure legends. A p value < 0.05 was considered statistically significant. Significance is indicated as follows: *p < 0.05, **p < 0.01, ***p < 0.001, ****p < 0.0001; “ns” indicates non-significant.

## Results

### Foxo3 is induced and accumulates in the nucleus of dopaminergic-like cells in response to mitochondrial and proteotoxic Parkinsonian stressors

Because mitochondrial dysfunction and aSyn aggregation represent two major and convergent pathogenic drivers in PD, we first asked whether Foxo3 responds to these insults in dopaminergic-like cells. We used SH-SY5Y cells as an *in vitro* surrogate, given their neuronal morphology and expression of key catecholaminergic markers, although they do not fully recapitulate mature dopaminergic neurons, and exposed them either to the mitochondrial complex I inhibitor MPP^+^ or to a truncated, aggregation-prone form of human α-synuclein (hSyn^TR110^).

Following 24 h of MPP^+^ exposure, SH-SY5Y cells displayed a dose-dependent increase in toxicity, as evidenced by increased detachment, pyknosis and cleaved caspase-3 staining (Fig. 1A–C). Strikingly, Foxo3 immunoreactivity increased in parallel and accumulated in the nucleus, with changes already detectable at low MPP^+^ concentrations. Foxo3 expression was increased after exposure to MPP^+^ at 5 µM (135% of control, p=0.0022; Fig. 1D), with concomitant increase of its nuclear localization (127% of control, p=0.016; Fig. 1E), preceding overt cell death, indicating that Foxo3 activation is an early stress response rather than a consequence of degeneration. Thus, Foxo3 is rapidly mobilized in response to mitochondrial stress in dopaminergic-like cells.

**Figure 1.**
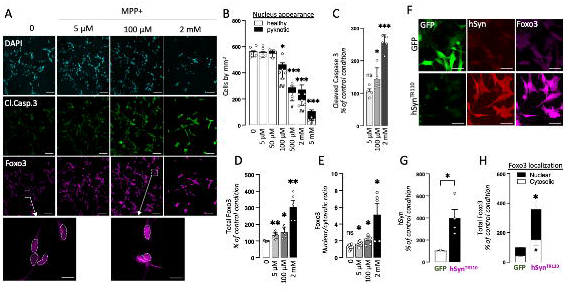
Foxo3 localization and apoptosis in SH-SY5Y cells following MPP^+^ treatment or α-synuclein overexpression. **(A)** Representative immunofluorescence images of SH-SY5Y cells treated with increasing concentrations of MPP^+^ for 24 h and stained for DAPI (cyan), cleaved caspase-3 (Cl. Casp3, green) and Foxo3 (magenta). Insets show higher magnification views with nuclei outlined based on DAPI staining. Scale bars: 100 µm (overview); 10 µm (insets). **(B)** Quantification of total, pyknotic and healthy cells across MPP^+^ concentrations. **(C)** Quantification of cleaved caspase-3 staining intensity per cell. **(D)** Quantification of total Foxo3 staining intensity per cell, normalized to untreated conditions. **(E)** Quantification of nuclear Foxo3 localization based on DAPI masking, expressed as nuclear-to-cytosolic ratio. **(F)** Representative immunofluorescence images of SH-SY5Y cells transduced with vectors expressing GFP (control) or hSynTR110 for 48 h and stained for GFP (green), hSyn (red) and Foxo3 (magenta). Scale bars: 20 µm. **(G)** Quantification of hSyn expression levels based on fluorescence intensity, normalized to control conditions. (H) Quantification of total and nuclear Foxo3 staining intensity following hSynTR110 expression, normalized to control conditions. **General:** Each dot represents one independent experiment (200–500 cells analyzed per condition); N = 6 independent experiments. **Statistics:** Wilcoxon test (comparison to reference condition). *p < 0.05; **p < 0.01; ***p < 0.001. In (B) and (H), # indicates comparison of pyknotic cells (B) or nuclear Foxo3 (H), whereas * indicates comparison of total cell number (B) or total Foxo3 staining (H).

We then asked whether a proteotoxic Parkinsonian insult would elicit a similar response. Viral overexpression of hSyn^TR110^ in SH-SY5Y cells resulted in robust intracellular α-synuclein accumulation (397.6% vs control, p=0.0267; Fig. 1F-G), and was associated with a marked increase in Foxo3 expression (357% of control, p=0.0174) and nuclear translocation (386%, p=0.0176; Fig. 1H).

Therefore, Foxo3 responds to both mitochondrial complex I inhibition and aSyn aggregation by increasing in abundance and relocating to the nucleus. Together, these data identify Foxo3 as a stress-responsive transcription factor activated by two central Parkinsonian insults, suggesting that it may participate in the coordination of downstream cellular responses beyond neurons. However, they do not indicate whether Foxo3 activation is adaptive or rather contributes to dopaminergic vulnerability.

### Constitutive Foxo3 deficiency confers robust neuroprotection across complementary models of Parkinsonian neurodegeneration

To determine whether Foxo3 functionally contributes to DNs degeneration in vivo, we analyzed constitutive Foxo3 knockout mice in two complementary mouse models of Parkinson’s disease.

We first used a model based on AAV-mediated unilateral intranigral expression of hSyn^TR110^, with GFP delivered to the contralateral side as an internal control (Fig. 2A-B; Additional file 1: Fig. S1). Eight weeks after induction, Foxo3^WT^ mice exhibited a marked reduction in TH^+^ neuronal cell bodies in the injected SNpc (35% loss), whereas Foxo3^KO^ mice showed only 16% loss, corresponding to 54% protection (p=0.0036; Fig. 2C-D). Consistently, TH^+^ fiber density in the dorsal striatum was reduced by 18% in Foxo3^WT^ mice but only 4% in Foxo3^KO^ mice (78% protection, p=0.0097; Fig. 2C,E).

**Figure 2.**
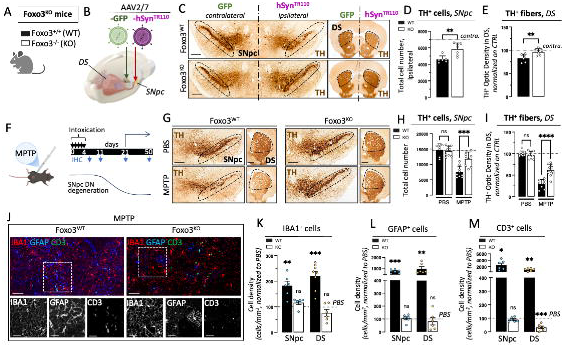
Foxo3 deficiency protects dopaminergic neurons in α-synuclein–driven and MPTP-induced models of Parkinson’s disease. **(A)** Schematic representation of Foxo3 knockout (KO) and wild-type (WT) mice used in the study. **(B)** Schematic representation of the experimental design for synucleinopathy induction by unilateral intranigral injection of AAV2/7-hSyn^TR110^, with contralateral injection of a control AAV-GFP vector. **(C)** Representative tyrosine hydroxylase (TH) immunostaining of the substantia nigra pars compacta (SNpc) and dorsal striatum (DS) eight weeks after AAV injection. Regions included in the analysis are outlined. Scale bars: 250 µm. **(D-E)** Quantification of TH^+^ neurons in the SNpc (D) and dopaminergic fiber density in the DS (E), normalized to the contralateral control side. Data are expressed as mean ± SEM. Each dot represents one mouse. Statistical analysis: Mann–Whitney test. **(F)** Schematic representation of the MPTP intoxication paradigm (daily intraperitoneal injections for five consecutive days). **(G)** Representative TH immunostaining of the SNpc and DS in WT and Foxo3 KO mice following MPTP treatment. Scale bars: 250 µm. **(H–I)** Quantification of TH^+^ neurons in the SNpc (H) and dopaminergic fiber density in the DS (I), expressed as percentage of PBS-treated controls. Data are expressed as mean ± SEM. Each dot represents one mouse. Statistical analysis: Two-way ANOVA with Tukey’s multiple comparisons test. **(J)** Representative immunostaining of microglia (IBA1), astrocytes (GFAP) and T cells (CD3) in the dorsal striatum. Insets show higher magnification. Scale bars: 100 µm (overview); 20 µm (insets). **(K–M)** Quantification of microglia (K), astrocytes (L) and T cells (M) density in the dorsal striatum, normalized to PBS-treated controls. Data are expressed as mean ± SEM. Each dot represents one mouse. Statistical analysis: Wilcoxon test.

We next assessed the impact of Foxo3 deficiency in a second, mechanistically distinct model based on systemic subacute MPTP intoxication (Fig. 2F). Dopaminergic neuron loss in this model becomes detectable from day 11 and stabilizes around day 21 (Additional file 2: Fig. S2A), enabling us to interrogate early molecular responses at day 10 and established neurodegeneration and neuroinflammation at day 21. In Foxo3^WT^ mice, MPTP induced a 48.2% loss of TH^+^ neurons in the SNpc, whereas Foxo3^KO^ mice exhibited only 18.8% loss, corresponding to 75% protection (p=0.0003; Fig. 2G-H). Axonal degeneration in the dorsal striatum was also significantly attenuated (70.8% vs 38.1% loss; p<0.0001; Fig. 2I). Importantly, this protective effect persisted over time, with 66.4% protection still observed at D50 (Additional file 2: Fig. S2B). Foxo3^WT^ and Foxo3^KO^ mice displayed comparable MPTP-to-MPP^+^ conversion, dopamine turnover (DOPAC/DA ratio), and DAT expression (Additional file 2: Fig. S2C–E), ruling out confounding differences in toxin handling or dopaminergic metabolism.

Both paradigms were associated, in Foxo3^WT^ mice, with a strong local glial and inflammatory response (Fig. 2J and Additional file 3: Fig. S3). MPTP treatment induced a marked increase in Iba1^+^ microglia (+182% SNpc, +224% DS), GFAP^+^ astrocytes (+682% SNpc, +1000% DS), and CD3^+^ T lymphocytes (+2494% SNpc, +1080% DS; Fig. 2K–M). Strikingly, these inflammatory changes were strongly reduced or absent in Foxo3^KO^ mice. Consistent with the quantification shown in Fig. 2M, CD3^+^ T cell density was significantly reduced following MPTP intoxication in Foxo3-deficient mice compared to their basal condition. Notably, however, these mice exhibited elevated T cell levels at baseline, such that post-MPTP values returned to levels comparable to WT controls under basal conditions. This bidirectional change suggests that Foxo3 deletion alters the baseline immune landscape and the dynamics of T cell responses to injury, rather than simply promoting or suppressing T cell accumulation. Comparable effects were observed in the hSyn^TR110^ model (Additional file 3: Fig. S3).

Together, these results demonstrate that constitutive Foxo3 deficiency confers robust and sustained protection against dopaminergic degeneration across complementary PD models, while markedly dampening associated neuroinflammatory responses. These findings identify Foxo3 as a key determinant of Parkinsonian vulnerability *in vivo* and raises the question of whether its pathogenic effects involve neuron-autonomous and/or extra-neuronal mechanisms.

### Foxo3 shapes early neuronal transcriptional responses to MPTP

To specifically assess the neuron-autonomous contribution of Foxo3 to dopaminergic neuron (DN) vulnerability, we generated mice in which Foxo3 was selectively deleted in DNs using DAT-driven Cre-mediated recombination (Foxo3^ΔDN^; Fig. 3A).

**Figure 3.**
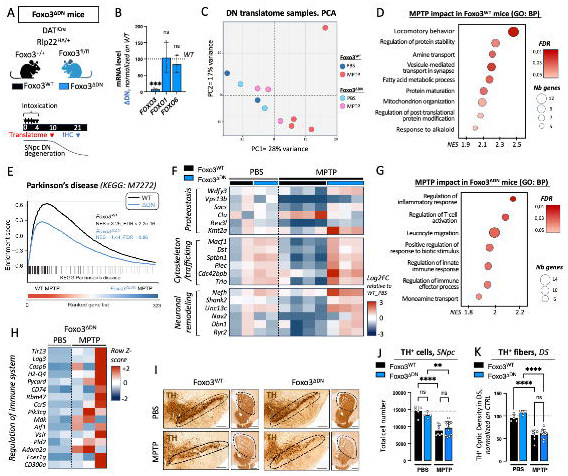
Neuron-specific deletion of Foxo3 attenuates early Parkinson-like transcriptional responses but does not confer neuroprotection. **(A)** Schematic representation of the experimental design for dopaminergic neuron-specific deletion of Foxo3 (Foxo3^ΔDN^) and MPTP intoxication paradigm. **(B)** qRT–PCR analysis of Foxo3, Foxo1 and Foxo6 expression in neuronal translatome samples from Foxo3^ΔDN^ mice, normalized to the mean level in Foxo3^WT^ samples. Each dot represents one sample (pools of 2 mice). Statistical analysis: Wilcoxon test. **(C)** Principal component analysis (PCA) of ribosome-associated mRNAs isolated from dopaminergic neurons (RiboTag) from Foxo3^ΔDN^ and Foxo3^WT^ mice following PBS or MPTP treatment. **(D)** Differential gene expression analysis between WT and Foxo3-deleted neurons following MPTP treatment. **(E)** GSEA of the KEGG Parkinson’s disease pathway in WT and Foxo3-deleted neurons following MPTP treatment. **(F)** Heatmap showing selected differentially expressed genes related to proteostasis, cytoskeletal organization and neuronal structural remodeling in Foxo3^WT^ and Foxo3^ΔDN^ dopaminergic neurons following PBS or MPTP treatment. **(G)** Dot plot showing immune-related pathways enriched in Foxo3^ΔDN^ dopaminergic neurons following MPTP treatment. (H) Heatmap showing row Z-scores of normalized expression values for immune-related genes in Foxo3^ΔDN^ neurons following PBS or MPTP treatments. Row Z-scores were calculated across all samples for each gene. **(I)** Representative tyrosine hydroxylase (TH) immunostaining of the substantia nigra pars compacta (SNpc) and dorsal striatum (DS) in Foxo3^ΔDN^ and Foxo3^WT^ mice following PBS or MPTP treatment. Scale bars: 250 µm. **(J-K)** Quantification of TH^+^ neurons in the SNpc (J) and dopaminergic fiber density in the striatum (K). Data are expressed as mean ± SEM. Each dot represents one mouse. Statistical analysis: two-way ANOVA with Tukey’s multiple comparison test.

We next investigated whether Foxo3 regulates early neuronal stress responses by profiling the dopaminergic translatome prior to overt degeneration using a DAT-driven RiboTag strategy (Additional file 4: Fig. S4A). Both Foxo3^WT^ and Foxo3^ΔDN^ mice were subjected to the MPTP paradigm, and ventral midbrain samples were collected at day 10 following intoxication, prior to overt neuronal loss (Additional file 2: Fig. S2A), enabling analysis of neuronal responses under conditions minimizing confounding inflammation. Quality controls confirmed robust enrichment of dopaminergic transcripts and efficient deletion of Foxo3 without compensatory changes in related family members (Fig. 3B, Additional file 4: Fig. S4B). Principal component analysis revealed a clear separation between PBS- and MPTP-treated Foxo3^WT^ samples, whereas Foxo3^ΔDN^ samples failed to segregate and instead clustered between WT-PBS and WT-MPTP groups (Fig. 3C), indicating a blunted transcriptional response to MPTP.

Differential expression analysis identified a substantial number of regulated genes in Foxo3^WT^ neurons following MPTP (Fold change >2 or <0.5, adjusted p < 0.05), supporting a robust transcriptional response. Gene Ontology enrichment analysis revealed coordinated regulation of pathways related to proteostasis, cytoskeletal organization, intracellular trafficking and neuronal structural remodeling in Foxo3^WT^ neurons following MPTP intoxication (Fig. 3D; Additional file 5: Fig. S5), indicating a robust activation of stress-adaptive and cellular remodeling programs. Notably, these pathways are hallmarks of Parkinson’s disease-associated neuronal dysfunction, including impaired proteostasis, cytoskeletal remodeling and altered intracellular trafficking, supporting the disease relevance of this transcriptional response. Consistent with this observation, gene set enrichment analysis revealed a strong induction of the KEGG Parkinson’s disease pathway in Foxo3^WT^ neurons following MPTP (NES = 3.25, FDR < 2.2×10^−16^), which was markedly attenuated in Foxo3^ΔDN^ neurons (NES = 1.44, FDR = 0.86; Fig. 3E). A gene-level view of this pathway further confirmed a coordinated induction of Parkinson’s disease-related genes in Foxo3^WT^ neurons that was broadly diminished in Foxo3^ΔDN^ neurons (Additional file 6: Fig. S6). A global view of differential gene expression between Foxo3^WT^ and Foxo3^ΔDN^ neurons under PBS and MPTP conditions is provided in Additional file 7: Fig. S7, highlighting the selective and context-dependent nature of transcriptional alterations associated with Foxo3 deletion.

Together, these findings indicate that MPTP induces a robust and disease-relevant transcriptional response in dopaminergic neurons, which is markedly attenuated in the absence of Foxo3.

### Foxo3-dependent neuronal responses involve cellular remodeling and immune-related pathways

We next examined the gene-level alterations underlying this transcriptional response to determine how Foxo3 shapes neuronal stress-adaptive programs. Analysis of selected differentially expressed genes revealed coordinated regulation of pathways related to proteostasis, cytoskeletal organization and neuronal structural remodeling (Fig. 3F). In Foxo3^WT^ neurons, MPTP induced a coherent modulation of genes involved in protein quality control and autophagy (e.g., *Wdfy3, Vps13b*, Clu), cytoskeletal organization and intracellular trafficking (e.g., *Macf1, Dst, Sptbn1, Cdc42bpb*), as well as synaptic and neuronal architecture (e.g., *Shank2, Unc13c, Nav2, Ryr2*). In contrast, this coordinated response was markedly altered in Foxo3^ΔDN^ neurons, where these pathways were no longer properly regulated and instead showed a blunted or partially opposite pattern of expression.

Strikingly, beyond this dysregulated stress response, Foxo3^ΔDN^ neurons exhibited a selective enrichment of immune-related pathways (Fig. 3G), indicating a shift toward neuroimmune-associated transcriptional programs. Gene-level analysis further revealed upregulation of immune-related transcripts, including *Lag3, H2-Q4, Cd74* and components of inflammatory signaling pathways (Fig. 3H), supporting the engagement of neuroimmune signaling in Foxo3-deficient neurons. Together, these findings suggest that loss of neuronal Foxo3 not only impairs adaptive stress programs but also promotes a transcriptional shift toward immune-associated pathways.

### Neuron-specific Foxo3 deletion does not confer neuroprotection

Despite these marked alterations in early neuronal transcriptional responses, Foxo3^ΔDN^ mice did not exhibit significant preservation of TH^+^ neurons in the SNpc or TH^+^ fibers in the dorsal striatum at D21 following MPTP intoxication (Fig. 3I-K), and no differences were observed compared to Cre-negative controls (Additional file 8: Fig. S8A-C). Similarly, neuroinflammatory parameters, including microgliosis, astrogliosis and T cell infiltration, were not detectably altered (Additional file 8: Fig. S8D-F). Importantly, a similar absence of neuroprotection was observed in the hSyn^TR110^ model (Additional file 9: Fig. S9), indicating that this lack of sustained neuroprotection is not restricted to the MPTP paradigm.

Hence, despite these early transcriptional alterations, Foxo3^ΔDN^ mice did not exhibit long-term neuroprotection, indicating that neuron-intrinsic Foxo3-dependent responses are insufficient to determine dopaminergic survival. These findings collectively indicate that while Foxo3 shapes early neuronal stress-adaptive programs, the dominant mechanisms governing dopaminergic vulnerability likely involve non-neuronal compartments, pointing to a central role for glial and immune cells.

### Microglia-specific deletion of Foxo3 is sufficient to protect dopaminergic neurons from MPTP-induced degeneration

Given that neuron-autonomous Foxo3 deletion failed to confer sustained protection, we next investigated whether dopaminergic vulnerability is determined by Foxo3-dependent mechanisms in non-neuronal compartments, with a particular focus on microglia. To this end, we generated inducible CX3CR1^CreERT2^;Foxo3^fl/fl^ mice, allowing selective deletion of Foxo3 in resident myeloid cells (Foxo3^ΔmG^; Fig. 4A). Efficient genetic deletion was evaluated at both the transcript and protein levels, showing marked reduction of Foxo3 expression in brain-resident myeloid cells without affecting peripheral myeloid compartments (Fig. 4B; Additional file 10: Fig. S10).

**Figure 4.**
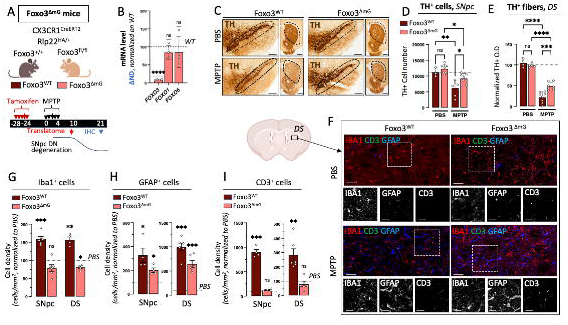
Microglial Foxo3 deletion confers neuroprotection and reshapes neuroinflammatory responses following MPTP intoxication. **(A)** Schematic representation of the experimental design for microglia-specific deletion of Foxo3 (Foxo3^ΔmG^) and MPTP intoxication paradigm. **(B)** qRT–PCR analysis of Foxo3, Foxo1 and Foxo6 expression in microglial translatome samples from Foxo3^ΔmG^ mice, confirming efficient and specific Foxo3 deletion. Expression levels are normalized to Foxo3^WT^ samples. Each dot represents one sample (pools of 2 mice). Statistical analysis: Wilcoxon test. **(C)** Representative tyrosine hydroxylase (TH) immunostaining of the substantia nigra pars compacta (SNpc) and dorsal striatum (DS) in Foxo3^ΔmG^ and Foxo3^WT^ mice following MPTP treatment. Scale bars: 250 µm. **(D–E)** Quantification of TH^+^ neurons in the SNpc (D) and dopaminergic fiber density in the striatum (E), normalized to PBS-treated controls. Each dot represents one mouse. Statistical analysis: two-way ANOVA with Tukey’s multiple comparisons test. **(F)** Representative confocal images of microglia (IBA1), astrocytes (GFAP) and T cells (CD3) in the dorsal striatum following MPTP treatment. Scale bars: 100 µm (overview); 20 µm (insets). **(G–I)** Quantification of microglia (G), astrocytes (H) and T cells (I), expressed as cell density and normalized to PBS-treated controls. Each dot represents one mouse. Statistical analysis: Wilcoxon test.

Following MPTP intoxication, Foxo3^ΔmG^ mice displayed robust preservation of TH^+^ dopaminergic neurons in the SNpc (25% vs 45% loss; p=0.0382; Fig. 4C-D) and striatal projections (50% vs 78% loss; p=0.0002; Fig. 4E). This level of neuroprotection was slightly lower than that observed in constitutive Foxo3-deficient mice (approximately 60% and 55% protection in the SNpc and striatum, respectively), but remained within a comparable range, indicating that microglial Foxo3 deletion accounts for a substantial fraction of the protective phenotype. In parallel, local glial and inflammatory responses were markedly attenuated, with reduced accumulation of Iba1^+^ microglia, GFAP^+^ astrocytes and CD3^+^ T cells in both the SNpc and dorsal striatum (Fig. 4F-I). Consistent with the quantification shown in Fig. 4F, Iba1^+^ microglial density was significantly reduced following MPTP intoxication in Foxo3^ΔmG^ mice compared to their basal condition. Notably, these mice exhibited elevated microglial density at baseline, such that post-MPTP levels approximated those observed in WT controls under basal conditions. This pattern suggests that Foxo3 deletion alters baseline microglial abundance and the dynamics of the response to injury, rather than simply reducing microglial activation.

Together, these findings demonstrate that microglia-specific deletion of Foxo3 is sufficient to confer substantial protection against MPTP-induced dopaminergic neurodegeneration.

### MPTP induces robust immune transcriptional remodeling in wild-type microglia

To identify microglial mechanisms associated with this protection, we profiled the microglial translatome using a RiboTag approach in the dorsal striatum and SNpc at day 10 following PBS or MPTP treatment (Experimental design: Fig. 4A). Efficient enrichment of microglial transcripts and depletion of neuronal markers were confirmed in immunoprecipitated samples (Additional file 11: Fig. S11A).

Principal component analysis revealed a clear segregation between WT_PBS and WT_MPTP samples, with PC1 accounting for 63% of the total variance, indicating a robust transcriptional response to MPTP in wild-type microglia (Fig. 5A). In Foxo3^WT^ microglia, MPTP exposure led to 868 upregulated and 457 downregulated genes in the dorsal striatum (adjusted p < 0.01), and similar changes in the SNpc (Additional file 11: Fig. S11B; Table 2), with detailed differential expression analyses for the SNpc provided in Additional file 12: Fig. S12. Pathway enrichment analysis revealed robust activation of both innate and adaptive immune signaling pathways (NES=1.95, FDR=0.0037 and NES=1.87, FDR=0.0107, respectively), including TLR and NF-κB cascades, as well as antigen processing and presentation pathways (Fig. 5B; Additional file 13: Fig. S13A).

**Figure 5.**
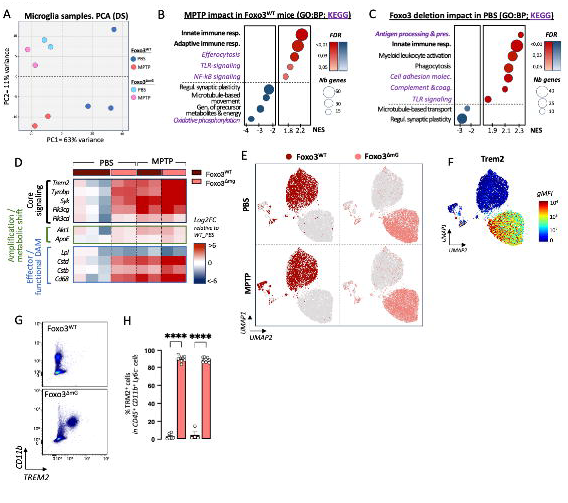
Microglial Foxo3 deletion establishes a distinct Trem2-associated basal state associated with selective immune remodeling. **(A)** Principal component analysis (PCA) of microglial translatome samples based on the 500 most highly expressed genes. **(B)** Dot plot showing pathways enriched in Foxo3^WT^ microglia following MPTP intoxication relative to PBS-treated controls. Dot size represents the number of genes associated with each pathway and color intensity indicates normalized enrichment scores (NES). **(C)** Dot plot showing pathways enriched in Foxo3^ΔmG^ microglia under basal conditions (PBS) relative to Foxo3^WT^ microglia. Dot size represents the number of genes associated with each pathway and color intensity indicates normalized NES. **(D)** Heatmap showing selected genes associated with Trem2/Tyrobp signaling (core and downstream pathways) in Foxo3^WT^ and Foxo3^ΔmG^ microglia following PBS or MPTP treatment. Gene expression values were normalized to the mean of WT_PBS samples and log2-transformed. **(E, F)** UMAP representation of high-dimensional flow cytometry analysis of microglia (CD45^+^ CD11b^+^ TMEM119^+^ Ly6C^-^ cells) from Foxo3^WT^ and Foxo3^ΔmG^ mice following PBS or MPTP treatment. (E) Distribution of cells from each experimental group across the global embedding. (F) Trem2 expression projected onto the shared UMAP space. **(G)** Representative flow cytometry density plots showing Trem2-expressing microglia in Foxo3^WT^ and Foxo3^ΔmG^ mice (PBS condition). **(H)** Quantification of Trem2^+^ cells in resident myeloid cells from Foxo3^WT^ and Foxo3^ΔmG^ mice. Data are expressed as mean ± SD. Each dot represents one mouse from 3 independent experiments. Statistical analysis: One-way ANOVA.

### Foxo3-deficient microglia display a pre-activated Trem2-associated immune state

Strikingly, the most pronounced transcriptional differences associated with Foxo3 deletion were already present under basal conditions rather than after MPTP exposure. Indeed, at steady-state, Foxo3^ΔmG^ microglia displayed extensive transcriptional remodeling, with 813 upregulated and 540 downregulated genes in the dorsal striatum as compared to Foxo3^WT^ microglia (Fold change > 2, adjusted p < 0.01; Additional file 13: Fig. S13B; Table 2). These changes reflected a coordinated enrichment of immune-related pathways (Fig. 5C and Additional file 13: Fig. S13A), including antigen processing and presentation (NES=2.34; FDR <2.2e^-16^), cell adhesion molecules (NES=2.12; FDR <2.2e^-16^), complement cascades (NES=2.10; FDR=0.0002), and Toll-like receptor signaling (NES=1.83; FDR=0.0259). Comparative analyses further revealed a partial overlap with previously described disease-associated microglia (DAM) signatures (Additional file 13: Fig. S13C), notably involving Trem2-associated and lysosomal programs, although several canonical DAM markers remained unchanged or only modestly affected. Despite this marked basal immune remodeling, no spontaneous dopaminergic alterations were observed (Fig 4D-E, left “PBS” parts), indicating that this baseline state is compatible with tissue homeostasis.

Notably, core components of the TREM2-TYROBP signaling module, including *Trem2, Tyrobp* and the downstream kinase *Syk*, were already elevated in Foxo3-deficient microglia under basal conditions, indicating that Foxo3 deletion intrinsically alters the baseline microglial state prior to neurotoxic challenge (Fig. 5D). This signature was associated with increased expression of lysosomal and phagocytic markers such as *Ctsd, Ctsb* and *Cd68*, whereas canonical lipid-associated DAM markers including *ApoE* and *Akt1* showed limited changes and *Lpl* expression was reduced, suggesting a selective remodeling of TREM2-associated microglial programs rather than a fully established DAM phenotype.

To further characterize microglial state remodeling at the protein level, we developed a high-dimensional 23-marker flow cytometry panel combining homeostatic, phagocytic, antigen-presentation and activation-associated microglial markers. High-dimensional flow cytometry analyses further revealed a marked redistribution of Foxo3^ΔmG^ microglia within the global UMAP space, largely associated with increased Trem2 expression (Fig. 5E-F). Importantly, TREM2 upregulation was observed across the overall Foxo3^ΔmG^ microglial population rather than within a discrete subcluster (Fig. 5G-H), supporting broad remodeling of microglial states.

Together, these findings indicate that Foxo3 deletion induces a selective and uncoupled activation of DAM-associated modules, characterized by enhanced Trem2–Tyrobp signaling and lysosomal/phagocytic programs in the absence of coordinated induction of lipid-associated pathways or loss of homeostatic microglial identity.

### Foxo3 deletion constrains canonical MPTP-induced microglial reprogramming

In contrast to Foxo3^WT^ microglia, which developed a distinct CD86^hi^/Clec7A^hi^ activated subpopulation following MPTP intoxication (Additional file 15: Fig. S15), Foxo3^ΔmG^ microglia did not exhibit expansion of this canonical MPTP-associated activation state, suggesting that Foxo3 deletion redirects microglial responses rather than globally suppressing activation. Consistent with this, Foxo3^ΔmG^ microglia displayed only limited additional transcriptional remodeling relative to baseline following MPTP exposure, as illustrated by the small number of differentially expressed genes between MPTP- and PBS-treated Foxo3^ΔmG^ microglia (Additional file 14: Fig. S14). Together with the PCA analysis (Fig. 5A), these findings indicate that Foxo3^ΔmG^ microglia remain transcriptionally close to their basal state after neurotoxic challenge, whereas Foxo3^WT^ microglia underwent extensive immune reprogramming involving innate and adaptive immune pathways, including TLR, NF-κB and antigen presentation programs.

Direct comparison between Foxo3^ΔmG^ and Foxo3^WT^ microglia following MPTP identified only 14 differentially expressed genes (adjusted p < 0.01; Additional file 11: Fig. S11C), indicating that neuroprotection is not associated with broad transcriptional reprogramming but rather with a constrained and selective response. Among these, Tyrobp remained significantly upregulated in Foxo3-deficient microglia, further supporting persistence of the Trem2-associated basal microglial state after injury. In addition, genes involved in innate immune signaling and complement regulation, including Wdfy1 and Csmd1, were differentially expressed, suggesting selective modulation of intracellular signaling pathways rather than generalized inflammatory activation.

Together, these findings identify Foxo3 as a regulator of microglial state-setting and indicate that baseline microglial configuration, rather than the magnitude of injury-induced activation, shapes dopaminergic vulnerability.

## Discussion

Our study identifies Foxo3 as a context-dependent regulator of dopaminergic vulnerability in experimental Parkinson’s disease and shows that its dominant pathogenic role is not neuron-autonomous. Although Foxo3 is mobilized in dopaminergic-like cells by mitochondrial complex I inhibition and α-synuclein aggregation, sustained neuroprotection *in vivo* is reproduced by microglial - but not dopaminergic neuron-specific - deletion. These data therefore shift the center of gravity of Foxo3 function in PD from an exclusively neuronal stress-response factor to a broader regulator of neuroimmune vulnerability. Importantly, this effect does not appear to be driven by global suppression of inflammatory activation following injury, but rather by qualitative differences in how microglial responses are configured and deployed.

This interpretation is biologically plausible in light of the established functions of Foxo factors in cellular quality control, metabolism, oxidative stress resistance and stress adaptation (Webb & Brunet, 2014). More specifically, Foxo3 occupies a distinctive position at the intersection of stress signaling and aging biology. Foxo3 is one of the most consistently replicated human longevity-associated genes, and its activity has been linked to resilience against metabolic and oxidative stress rather than to simple suppression of cellular responses (Flachsbart et al., 2009; Willcox et al., 2008). This apparent paradox likely reflects the context-dependent nature of Foxo3 signaling, whose effects depend on cell type, activation state and disease context rather than being uniformly protective.

Aging can be viewed as a progressive shift in stress and immune set points, often referred to as inflammaging (Franceschi et al., 2018). In the brain, microglia are among the cell types most profoundly remodeled by age, acquiring altered inflammatory thresholds, phagocytic properties and interactions with adaptive immune cells (Deczkowska et al., 2018). Consistent with the idea that aging reshapes immune and stress-response thresholds, FOXO3 polymorphisms have been associated not only with longevity but also with modulation of inflammatory tone and disease severity in humans (Lee et al., 2013; Torigoe et al., 2024). In this context, our findings suggest that Foxo3 contributes to setting microglial stress and immune thresholds, thereby influencing how the aging brain converts neuronal stress into chronic neuroinflammation and degeneration.

The neuronal data nonetheless remain informative. In dopaminergic neurons, Foxo3 deletion attenuated the early translational response to MPTP, including a marked reduction in enrichment of the KEGG Parkinson’s disease pathway. This indicates that Foxo3 contributes to the acute molecular response mounted by neurons facing mitochondrial intoxication. Yet this buffering of the early neuronal Parkinson-like signature did not translate into improved long-term survival. The dissociation between early molecular attenuation and unchanged degeneration suggests that Foxo3-dependent neuronal stress programs are not the principal determinants of dopaminergic fate in vivo, but rather represent an early adaptation to injury that is insufficient to redirect disease outcome. In other words, neuronal Foxo3 shapes how dopaminergic neurons initially register the insult, but not whether they ultimately survive it. Importantly, this dissociation indicates that attenuation of early Parkinson-like molecular signatures is not sufficient to alter long-term disease trajectory *in vivo*.

An additional clue comes from the emergence of immune-related transcripts in Foxo3-deficient neurons after MPTP. Although this program was limited, it is conceptually important. Neurons can express immune-associated molecules under stress, including MHC class I components, and catecholaminergic neurons seem particularly vulnerable to immune recognition mechanisms. In Parkinsonian contexts, neuronal MHC-I expression has been linked to susceptibility to T-cell-mediated injury, while α-synuclein-specific T cell responses have been demonstrated in patients (Cebrián et al., 2014; Sulzer et al., 2017). More broadly, neuronal expression of MHC-I has been implicated in synaptic plasticity and activity-dependent remodeling (Huh et al., 2000; Shatz, 2009). In that light, the immune-like shift observed in Foxo3^ΔDN^ neurons may reflect altered neuroimmune communication rather than protective adaptation. This provides a plausible bridge between the attenuated neuronal Parkinson signature and the persistent degeneration observed in Foxo3^ΔDN^ mice: even when part of the canonical neuronal stress program is blunted, the inflammatory microenvironment may continue to impose sufficient extrinsic pressure to drive degeneration.

In contrast, microglia-specific deletion of Foxo3 was sufficient to reproduce a substantial part of the constitutive knockout phenotype, including preservation of nigral cell bodies, striatal innervation and attenuation of neuroinflammatory hallmarks. This identifies resident myeloid cells as key drivers of the protective phenotype. However, the magnitude of protection remained moderately reduced compared to constitutive Foxo3 knockout mice, suggesting that additional cell types may contribute to the full phenotype. Importantly, this does not challenge the central role of microglia but rather refines it, suggesting that microglial Foxo3 acts as a primary driver within a broader multicellular context. Accordingly, these findings support the now well-established view that microglial state transitions are not merely epiphenomena of neuronal injury but active regulators of disease progression (Hickman et al., 2018; Keren-Shaul et al., 2017; Masuda et al., 2019).

Together, these observations are consistent with a broader role of Foxo3 in macrophage biology beyond the CNS. In peripheral macrophages, Foxo3 has been implicated in the regulation of inflammatory tone, cytokine production and stress adaptation, notably through modulation of NF-κB-dependent pathways and control of pro-inflammatory cytokine expression (Dejean et al., 2009; Lin et al., 2004). This raises the possibility that Foxo3 may act more broadly across tissue-resident macrophage populations rather than as a strictly microglia-specific regulator. An important consideration is that CX3CR1-driven targeting is not restricted to parenchymal microglia and could also affect other CNS-resident macrophage populations, including border-associated macrophages (BAMs). Recent studies have demonstrated that BAMs actively contribute to neuroinflammatory responses in α-synuclein-based models of Parkinson’s disease, particularly through antigen presentation and regulation of immune cell recruitment (Schonhoff et al., 2023). More generally, transcriptomic and functional analyses have highlighted the distinct identity and roles of CNS-associated macrophage populations, including BAMs, at brain interfaces (Brioschi et al., 2021; Van Hove et al., 2019), which originate from embryonic precursors distinct from circulating monocytes (Goldmann et al., 2016). These findings suggest that CNS macrophage populations beyond microglia may participate in shaping neurodegenerative trajectories. Future work will therefore be required to disentangle the respective contributions of microglia and BAMs to the protective phenotype observed here.

A central finding of this study is that Foxo3 does not primarily control the magnitude of the microglial response following injury, but instead shapes the baseline configuration from which this response unfolds. The most prominent transcriptional effects of Foxo3 deletion were already evident under homeostatic conditions, establishing a distinct microglial state that is maintained without compromising nigrostriatal integrity at rest. Following MPTP intoxication, Foxo3-deficient microglia exhibited only limited additional transcriptional reprogramming, indicating that the protective phenotype does not arise from suppression of activation but from qualitative reshaping of response trajectories. In parallel, high-dimensional cytometry analyses revealed that WT microglia developed a distinct CD86^hi^/Clec7A^hi^ activated subpopulation after MPTP exposure that was not similarly expanded in Foxo3^ΔmG^ microglia. Thus, Foxo3 deletion appears to redirect microglial state transitions away from a canonical MPTP-induced activation trajectory rather than broadly suppressing immune responsiveness. In this context, Foxo3 functions less as a binary regulator of activation than as a rheostat shaping how microglial responses are configured and amplified during neurodegenerative stress.

The widespread upregulation of TREM2 across microglia in Foxo3^ΔmG^ mice is particularly notable. TREM2-TYROBP signaling is a central regulator of adaptive microglial responses to neurodegenerative stress, including damage sensing, metabolic adaptation and containment of pathology, and is required for effective microglial engagement with neurodegenerative damage (Keren-Shaul et al., 2017; Ulland & Colonna, 2018; Wang et al., 2015). More recent studies further emphasize that TREM2 signaling is highly context-, stage- and region-dependent, which may account for apparent discrepancies across models of neurodegeneration (Alonge et al., 2025; Hou et al., 2026).

Our data are consistent with this evolving view but provide an important mechanistic extension. Foxo3 deletion was associated with coordinated reinforcement of core components of the TREM2-TYROBP signaling module already at baseline, together with activation of downstream lysosomal and signaling pathways. Importantly, this state did not correspond to full acquisition of a canonical DAM phenotype, but rather to selective engagement of Trem2-associated programs within a broader immune-associated basal microglial configuration. This distinction is important because most studies have focused on microglial responses after neurodegenerative stimuli, whereas our data indicate that a predefined basal microglial state can shape subsequent disease trajectory. Interestingly, components of the TREM2-TYROBP module and associated pathways identified in our dataset overlap with gene programs linked to IL-10-mediated regulation, suggesting partial convergence between damage-sensing, metabolic adaptation and regulatory microglial programs. Consistent with this idea, microglia-specific IL-10 expression was recently shown to limit neuroinflammation and dopaminergic degeneration in Parkinsonian models (Bido et al., 2024). Thus, Foxo3 deletion may bias microglia toward protective signaling programs without broadly suppressing immune responsiveness. While these observations support a role for TREM2-associated pathways in the protective phenotype, direct causal relationships remain to be established.

At the intracellular level, this reconfiguration is further supported by selective rewiring of signaling pathways. In particular, modulation of Wdfy1 suggests that protection arises from selective adjustment of intracellular signaling dynamics rather than broad suppression of immune activation. Wdfy1 has been implicated in endosomal TLR3/TLR4–TRIF signaling and amplification of innate immune responses (Hu et al., 2015), and its downregulation is consistent with attenuation of intracellular amplification loops rather than suppression of activation per se. More broadly, these observations are compatible with emerging concepts of innate immune training, whereby prior microglial states durably shape subsequent responses to injury (Netea et al., 2020; Wendeln et al., 2018). In our model, Foxo3 deletion appears to establish a preconfigured microglial state with restrained injury-induced reprogramming, allowing microglia to respond to stress without transitioning into maladaptive inflammatory amplification.

These findings support a compartment-specific model of Foxo3 function in neurodegeneration. In neurons, Foxo3 contributes to acute adaptation to mitochondrial stress, whereas in microglia it sets signaling thresholds and response trajectories that become pathogenic in the context of chronic neurodegenerative stress. If Foxo3 indeed acts as a resilience-associated gene in humans (Morris et al., 2015), its effects are therefore likely to depend strongly on cellular context and tissue compartment rather than being uniformly protective.

From a therapeutic perspective, our findings suggest that targeting FOXO3-dependent pathways may offer an alternative strategy to conventional anti-inflammatory approaches by resetting microglial response thresholds rather than suppressing activation per se. Such interventions could promote protective microglial states without compromising essential immune functions, consistent with emerging concepts that emphasize the role of microglial state modulation in neurodegenerative diseases (Deczkowska et al., 2018; Hickman et al., 2018). However, given the pleiotropic and context-dependent roles of FOXO3 across cell types, achieving sufficient cell-type specificity will be a major challenge for future translational applications. Importantly, public single-cell transcriptomic datasets from Parkinson’s disease brains indicate that FOXO3 expression is broadly maintained across human microglial populations, in contrast to canonical DAM-associated genes that remain restricted to activated subsets (Additional file 16: Fig. S16; Martirosyan et al., 2024). These observations are consistent with the idea that Foxo3 acts upstream of global microglial state regulation rather than marking a specific activated subset.

Taken together, our findings redefine Foxo3 as a regulator of microglial state-setting rather than a purely neuronal stress-response factor, and identify baseline microglial configuration as a key determinant of neurodegenerative trajectory. More broadly, these findings position microglial state-setting as a central organizing principle of neurodegeneration, with implications extending beyond Parkinson’s disease to brain aging and neuroimmune dysfunction.

## Acknowledgments

We thank Dr. Ludovic Tricoire (P. Faure’s laboratory, ESPCI, Paris, France) for providing the DAT-iCre mouse line. Flow cytometry and imaging experiments were performed at the Univ Toulouse, INSERM, CNRS, Infinity core facility connected to Toulouse Réseau Imagerie network, member of the France-BioImaging national infrastructure supported by the French National Research Agency (ANR-24-INBS-0005 FBI BIOGEN). We thank Simon Lachambre for assistance with imaging experiments. We also thank Adeline Chaubet for support with transcriptomic sample preparation, and the GeT platform (Genotoul, Toulouse) as well as the animal facility CREFRE (Inserm US006) for technical support and animal care.

We acknowledge Prs. Steven’s and Blom’s laboratories (CMSD1 consortium) for providing reagents and valuable input, even though antibody validation for IHC with Iba1 was not successful in our hands.

We thank Dr. Hélène Hirbec for insightful discussions on microglial signatures, and Pr. Roland Liblau for critical reading of the manuscript.

## Author contributions

Conceptualization: M.S., A.S.D.

Methodology: M.S., B.T., S.R., Q.H.

Investigation: B.T., S.R., Q.H., M.P., M.A., L.D.-C., D.F., M.Z., A.-L.L.D., A.-L.I.

Formal analysis: M.S., M.Z., B.T., S.R.

Resources: D.A.-F., B.K., S.H., S.F.

Data curation: M.Z.

Writing - original draft: M.S.

Writing - review & editing: M.S., S.F.

Supervision: M.S.

Funding acquisition: M.S.

