## Supplementary figures and images for "Microglial Foxo3 shapes dopaminergic vulnerability in Parkinson’s disease"

### SupFig1.png

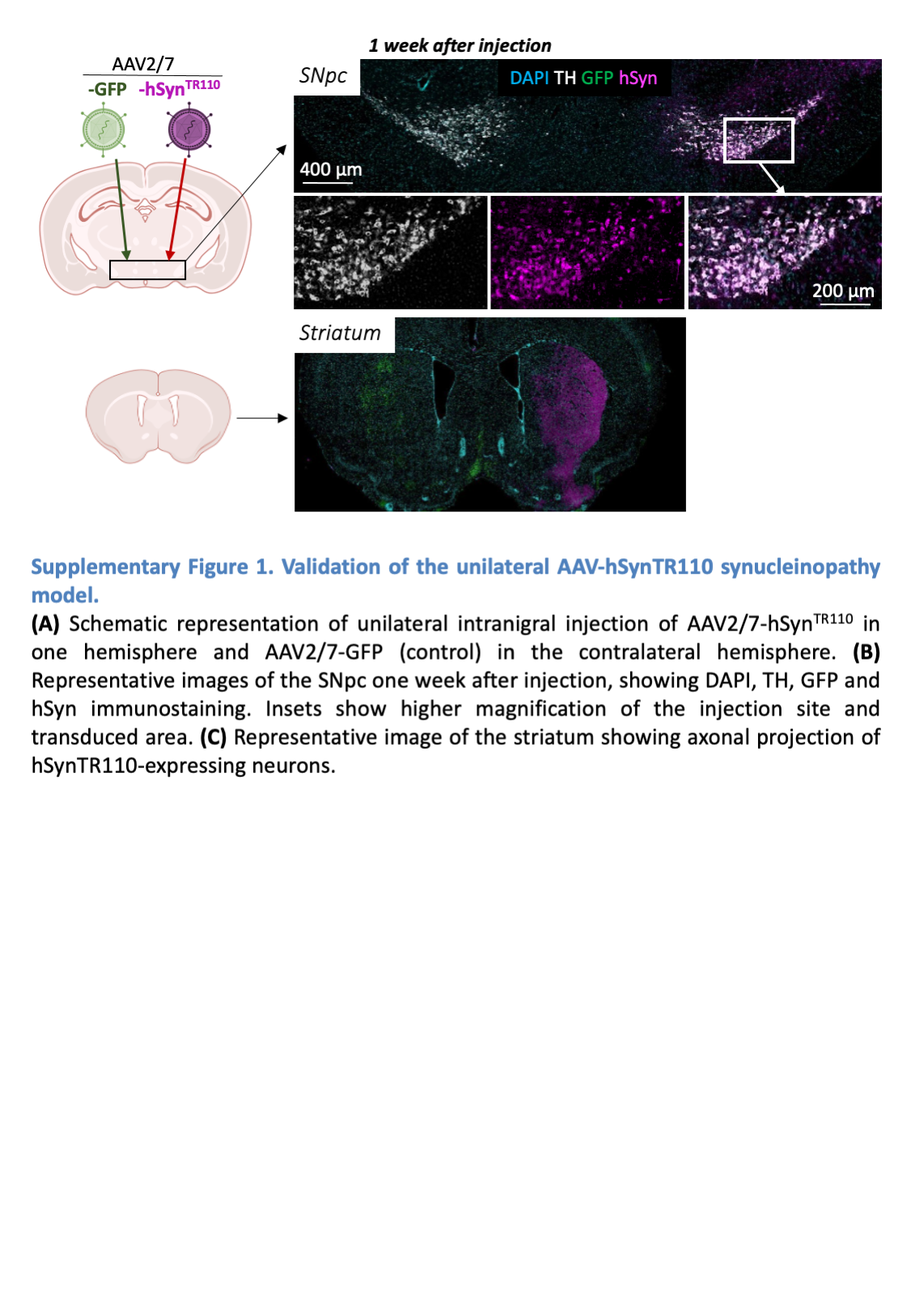

### SupFig2.png

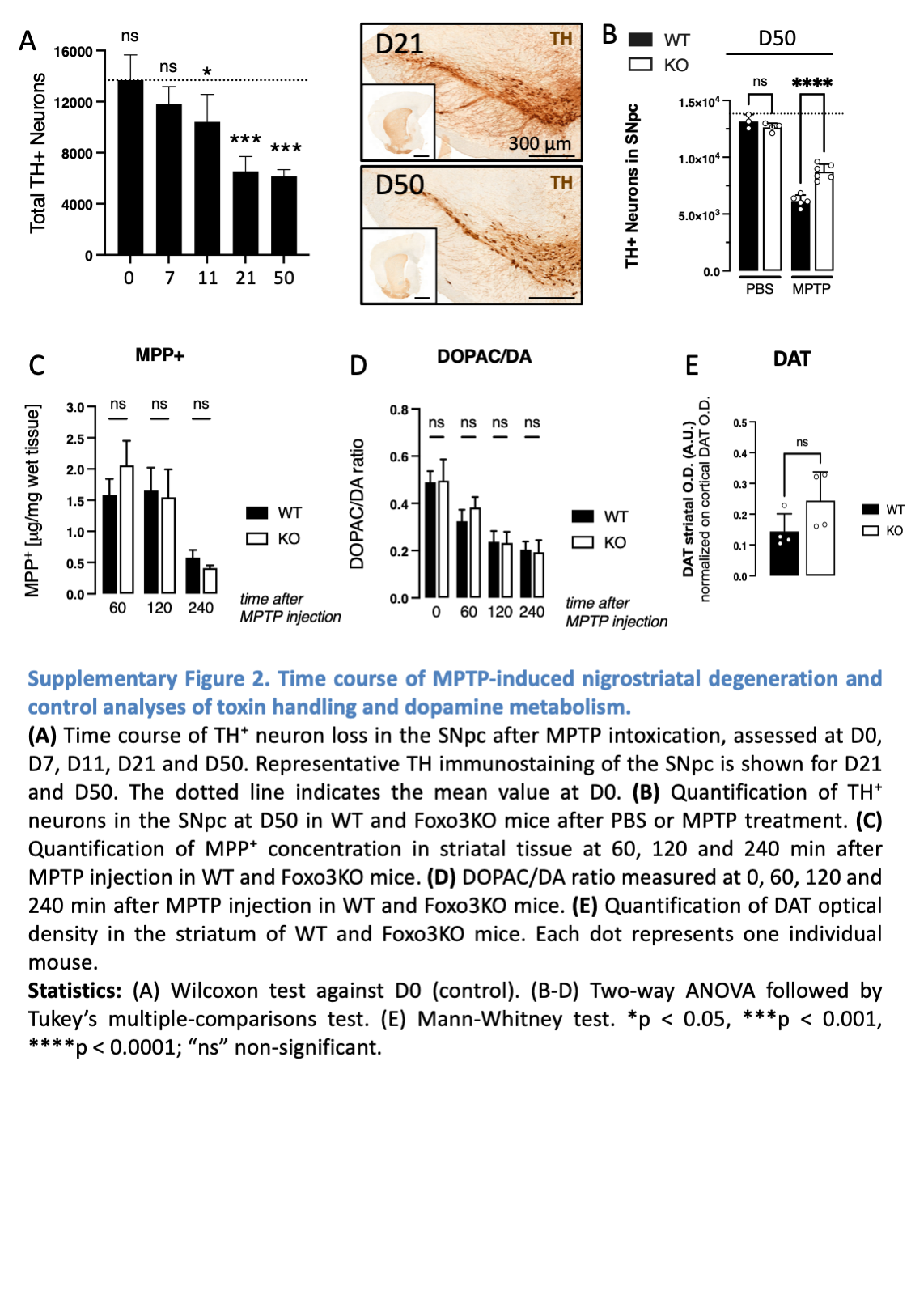

### SupFig3.png

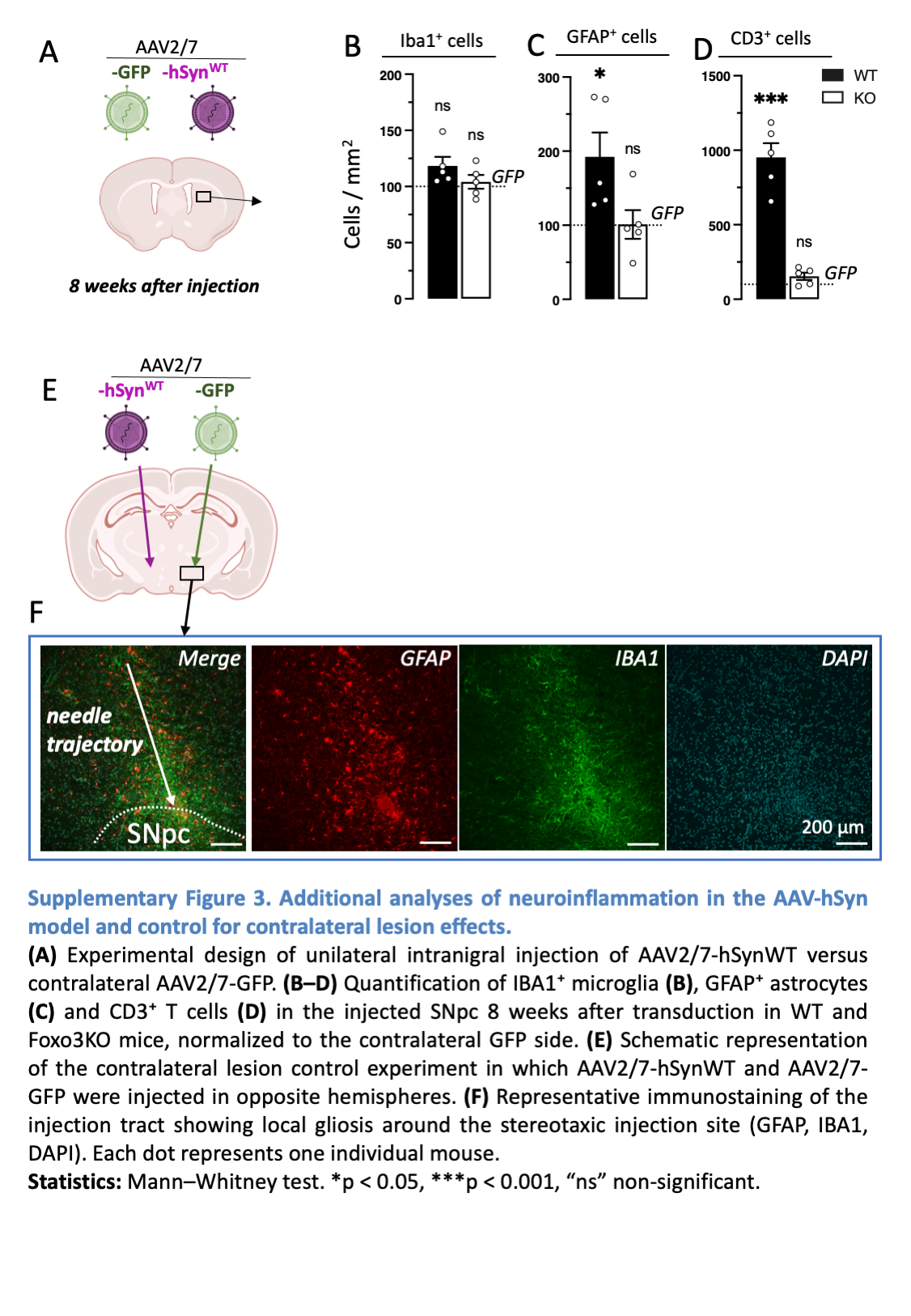

### SupFig4.png

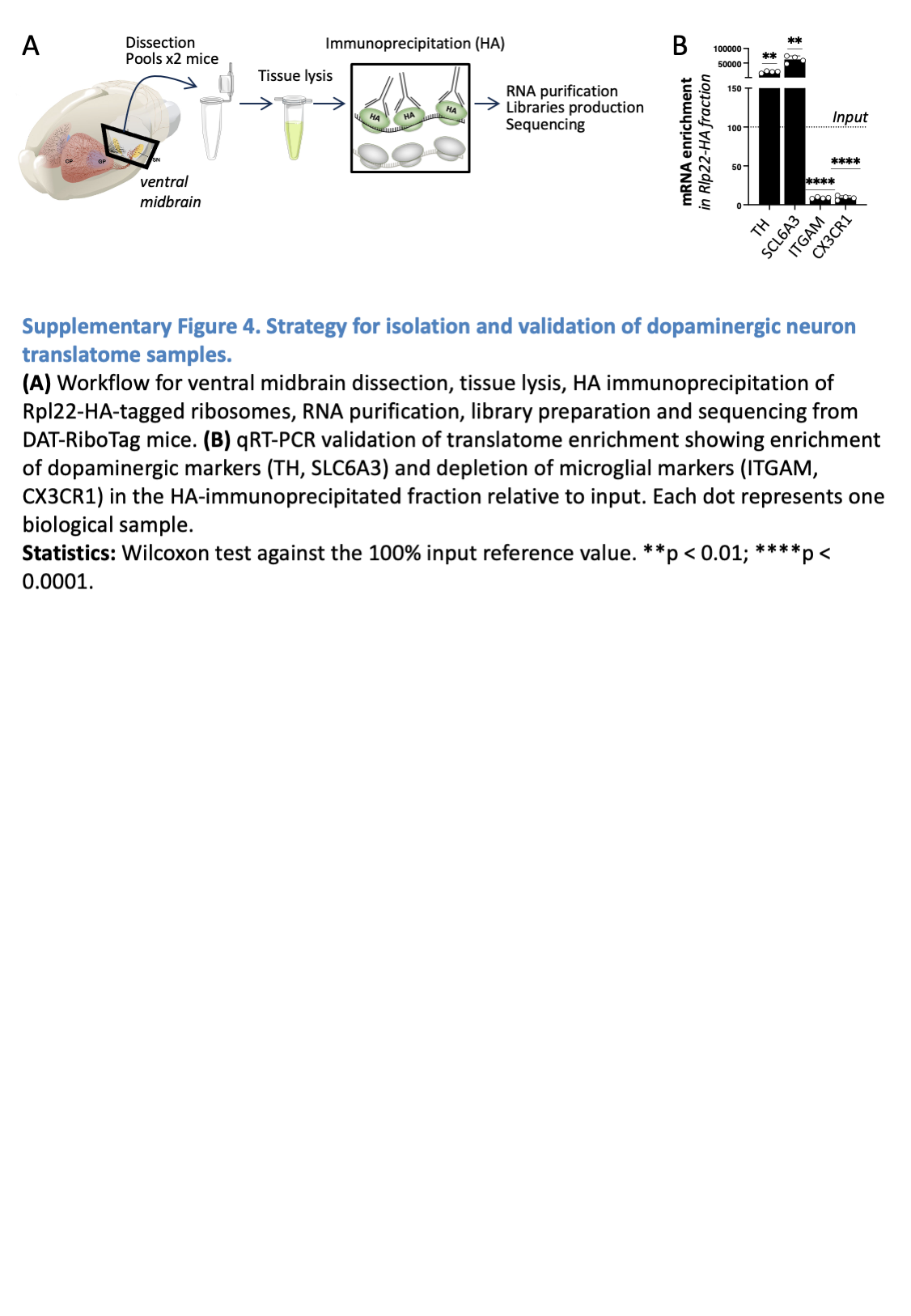

### SupFig5.png

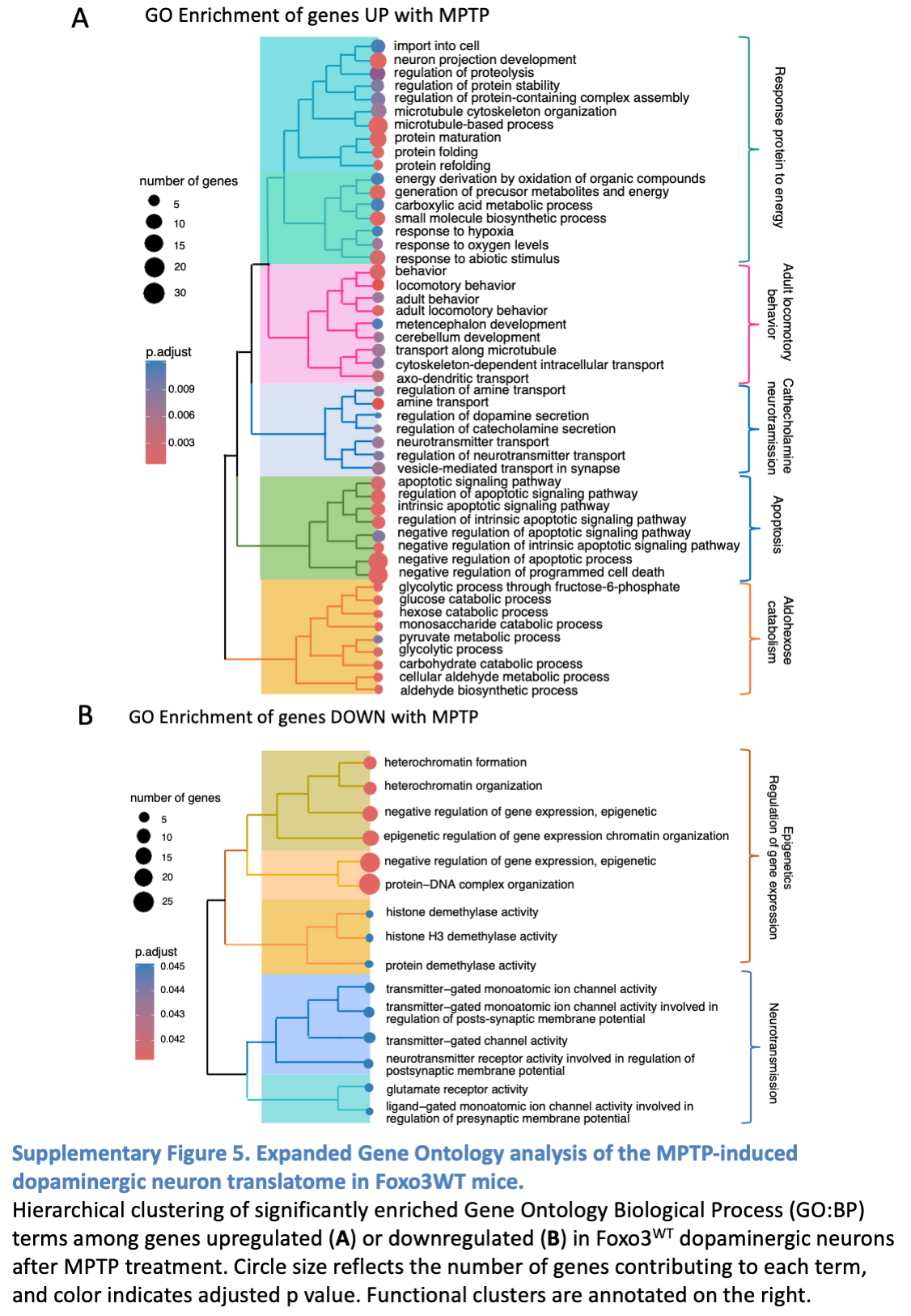

### SupFig6.png

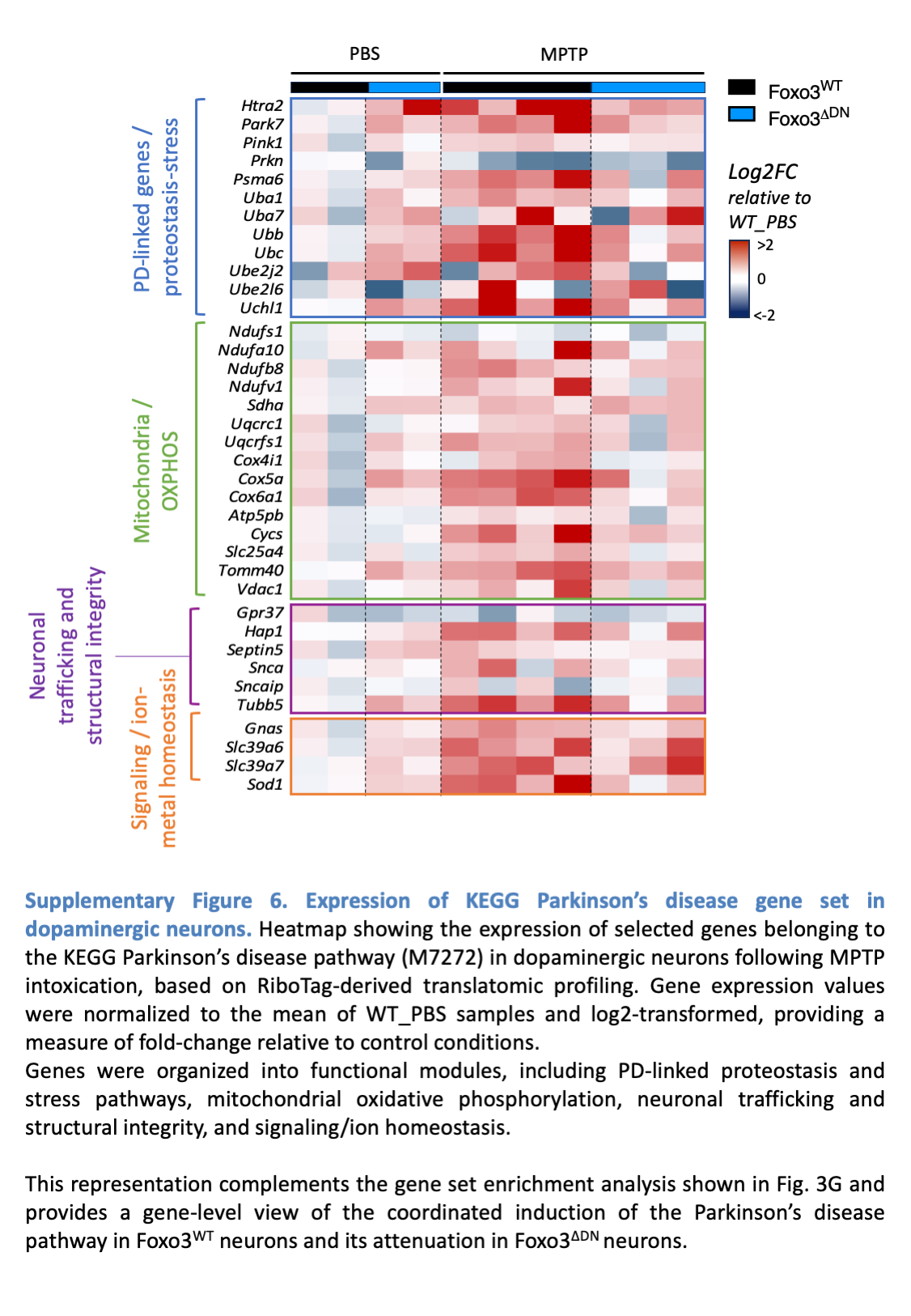

### SupFig7.png

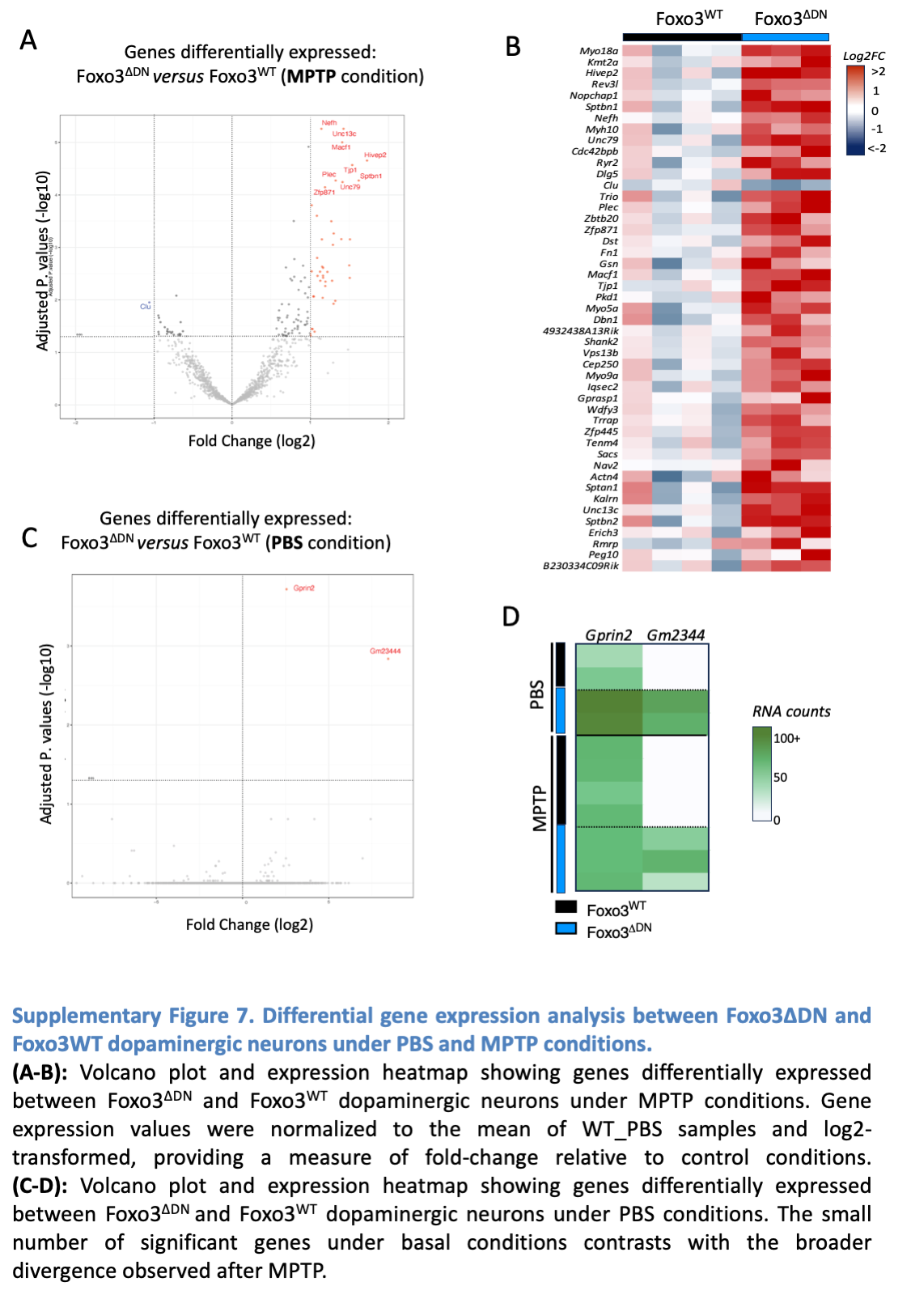

### SupFig8.png

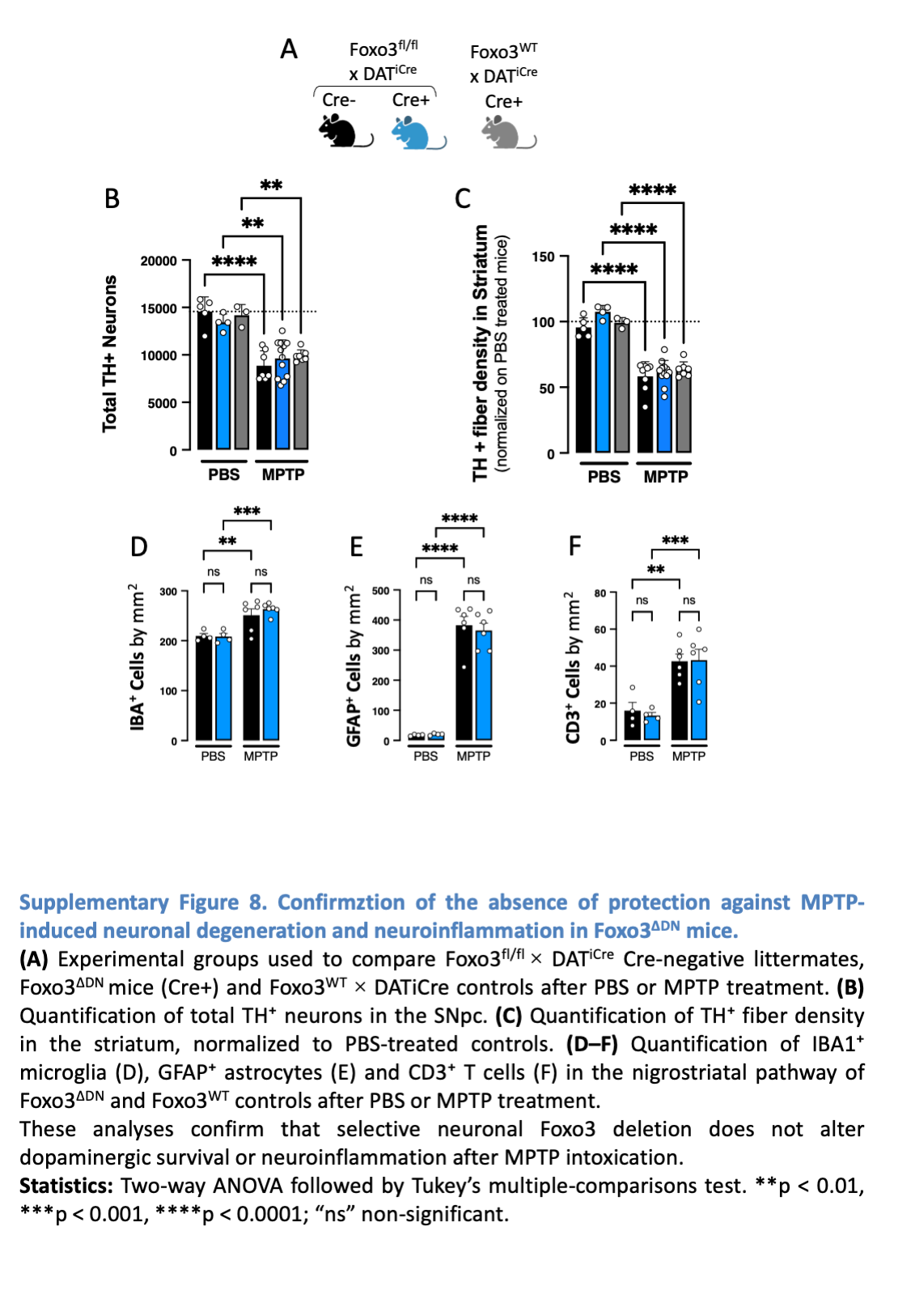

### SupFig9.png

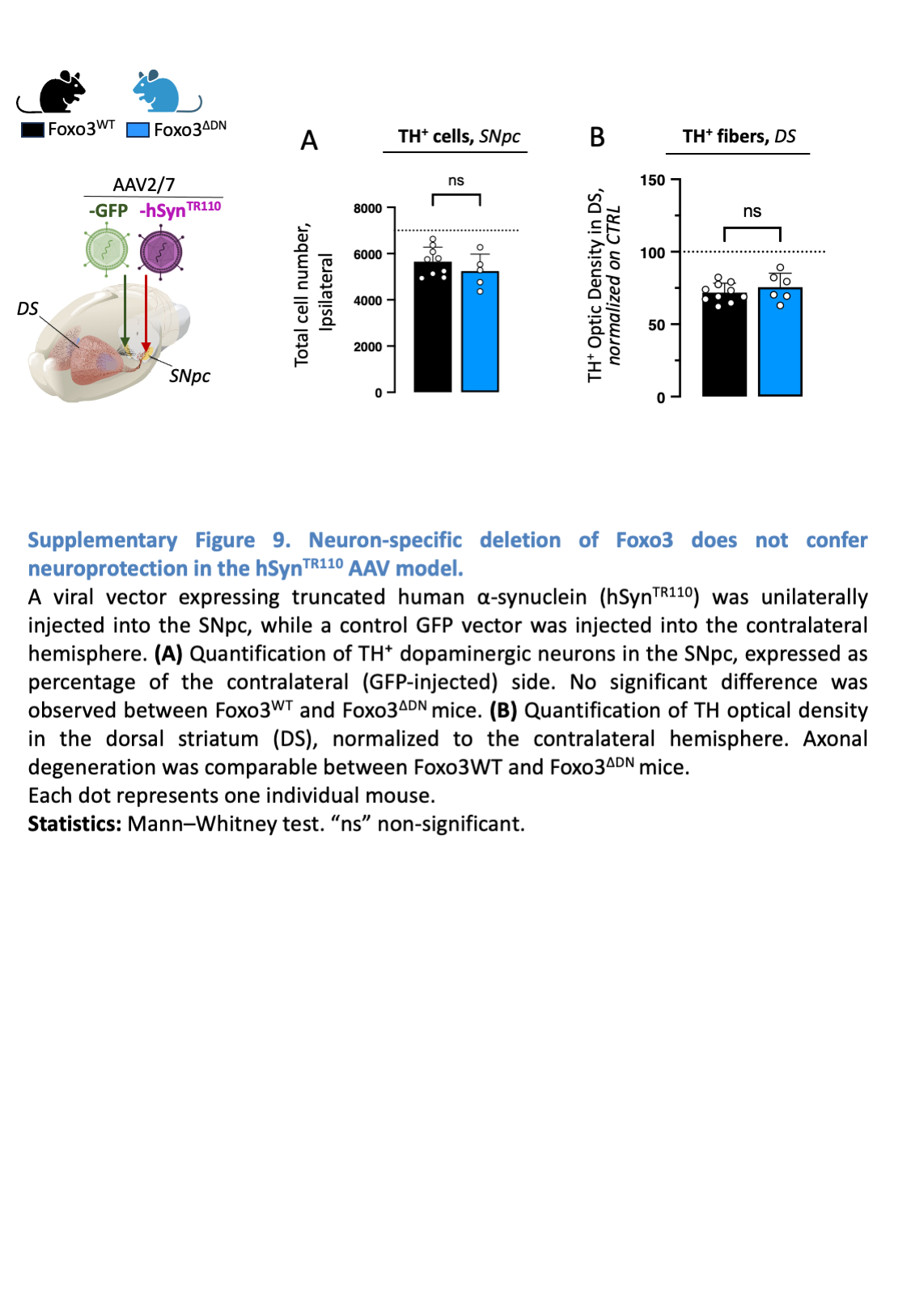

### SupFig10.png

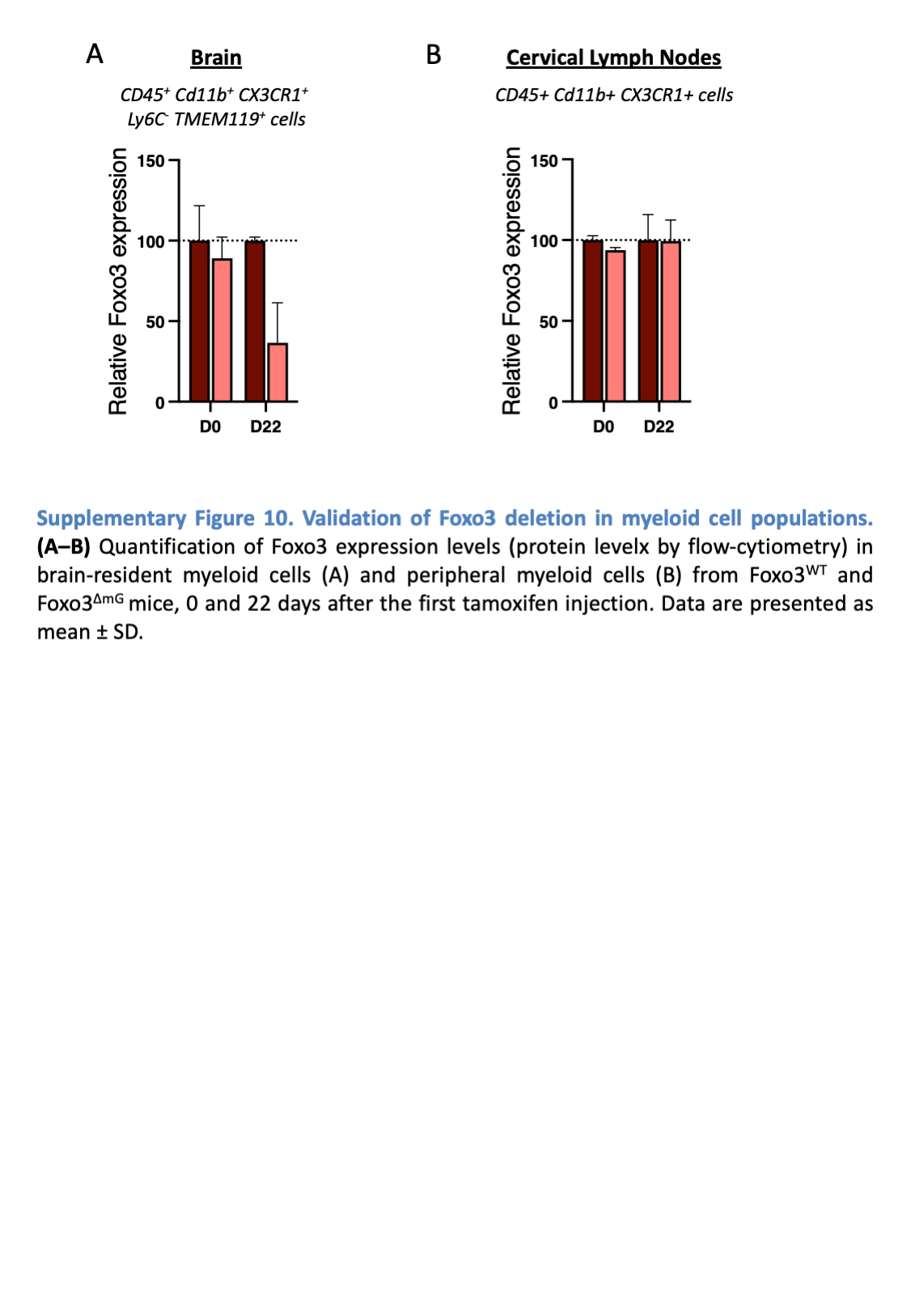

### SupFig11.png

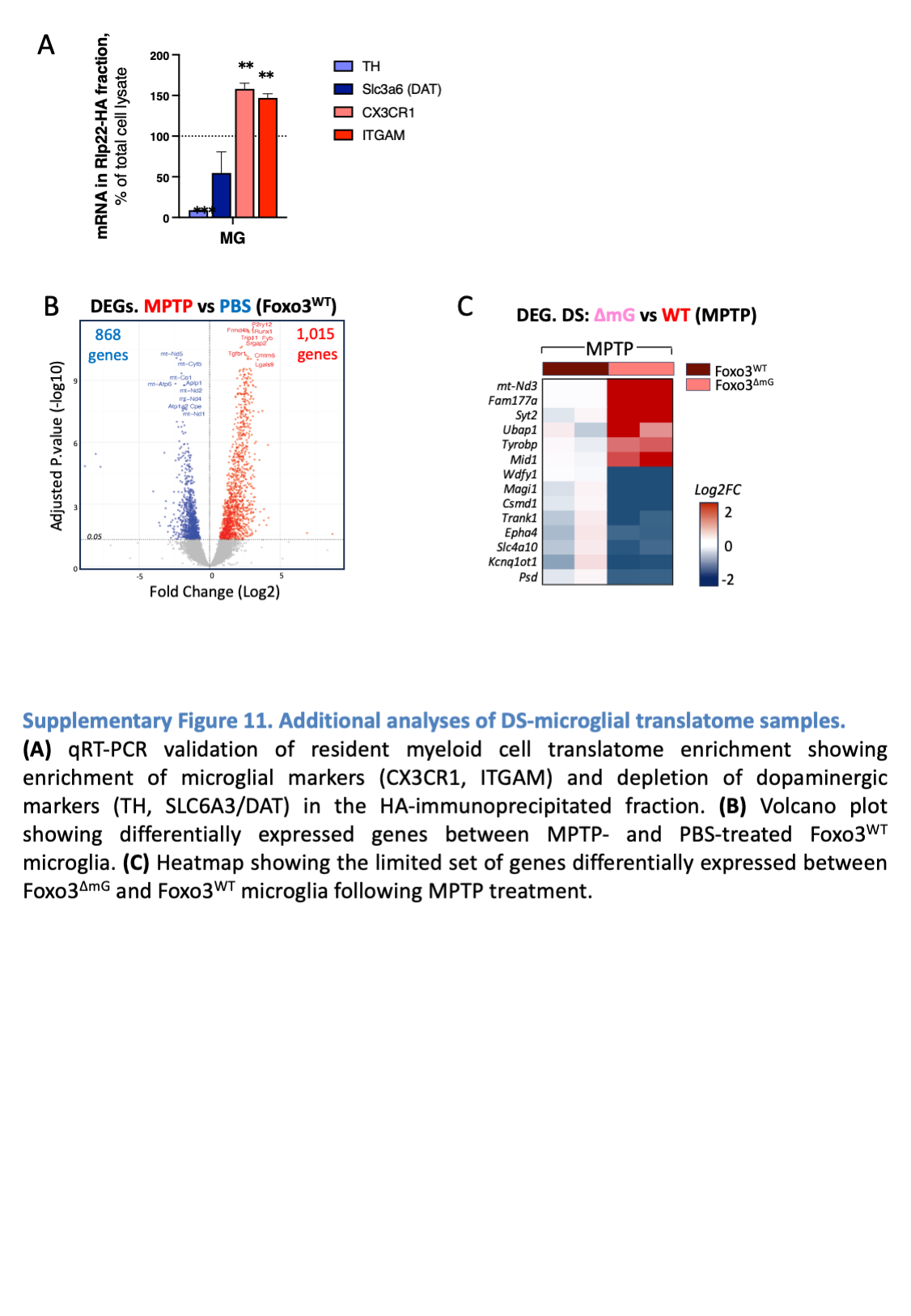

### SupFig12.png

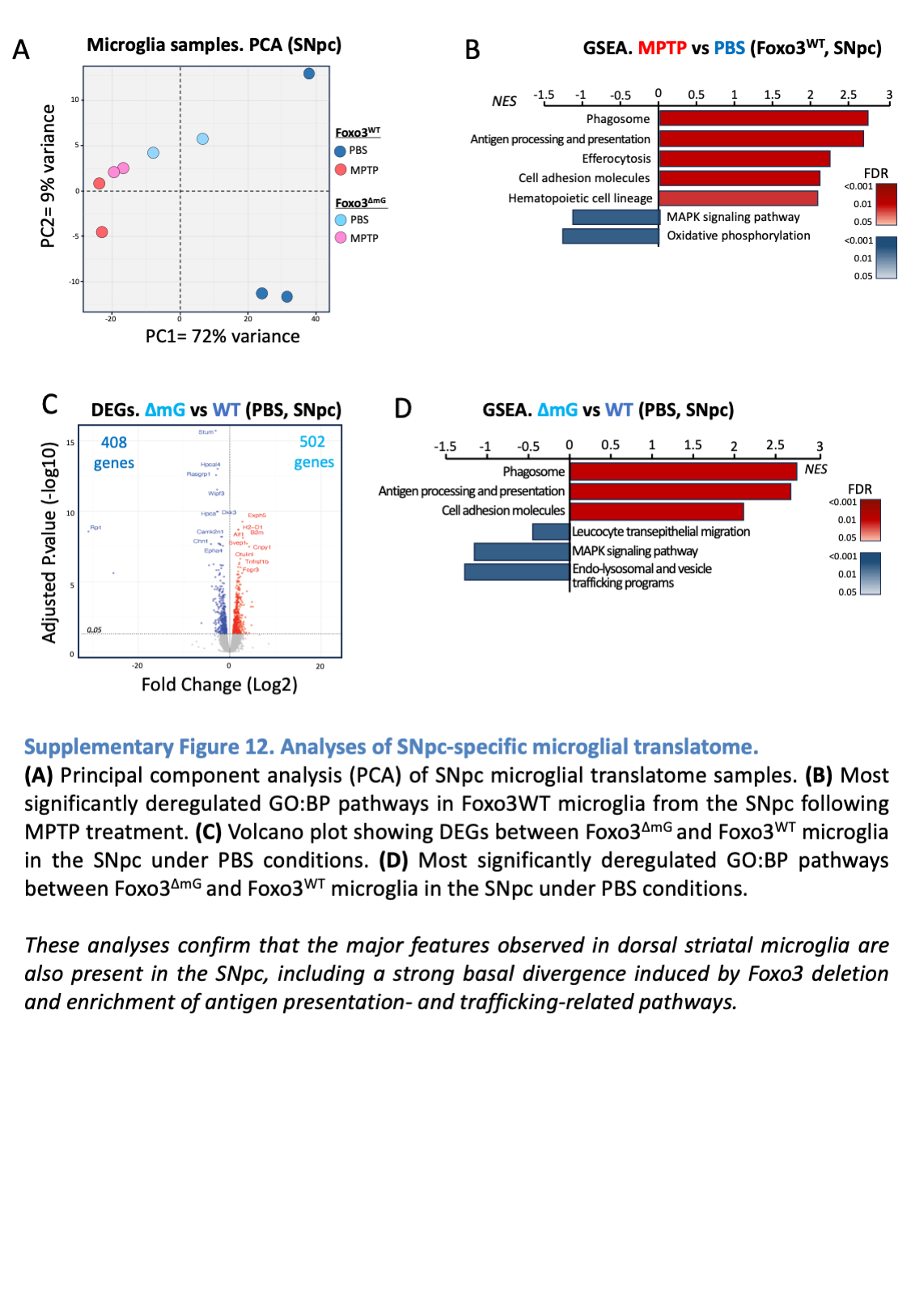

### SupFig13.png

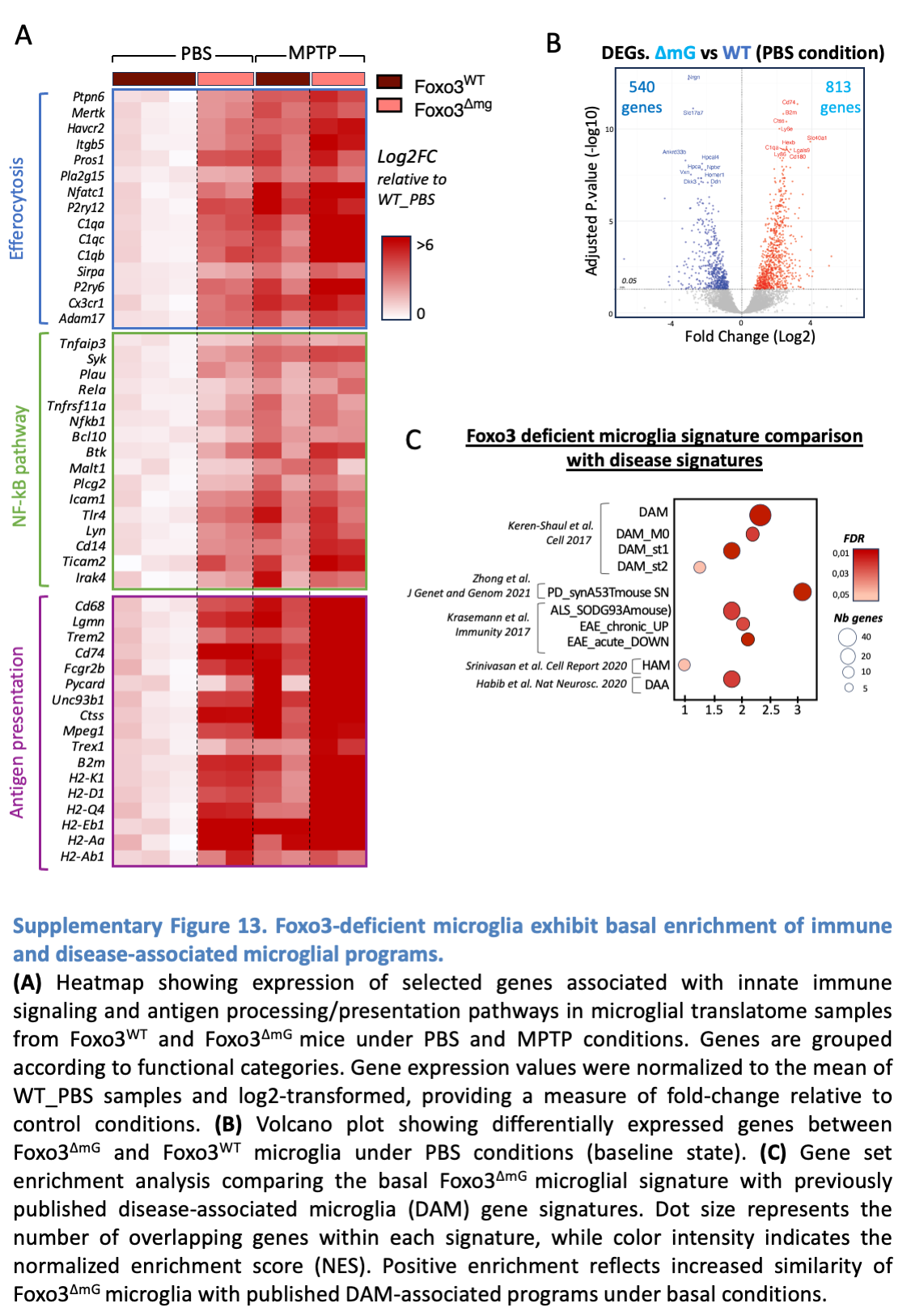

### SupFig14.png

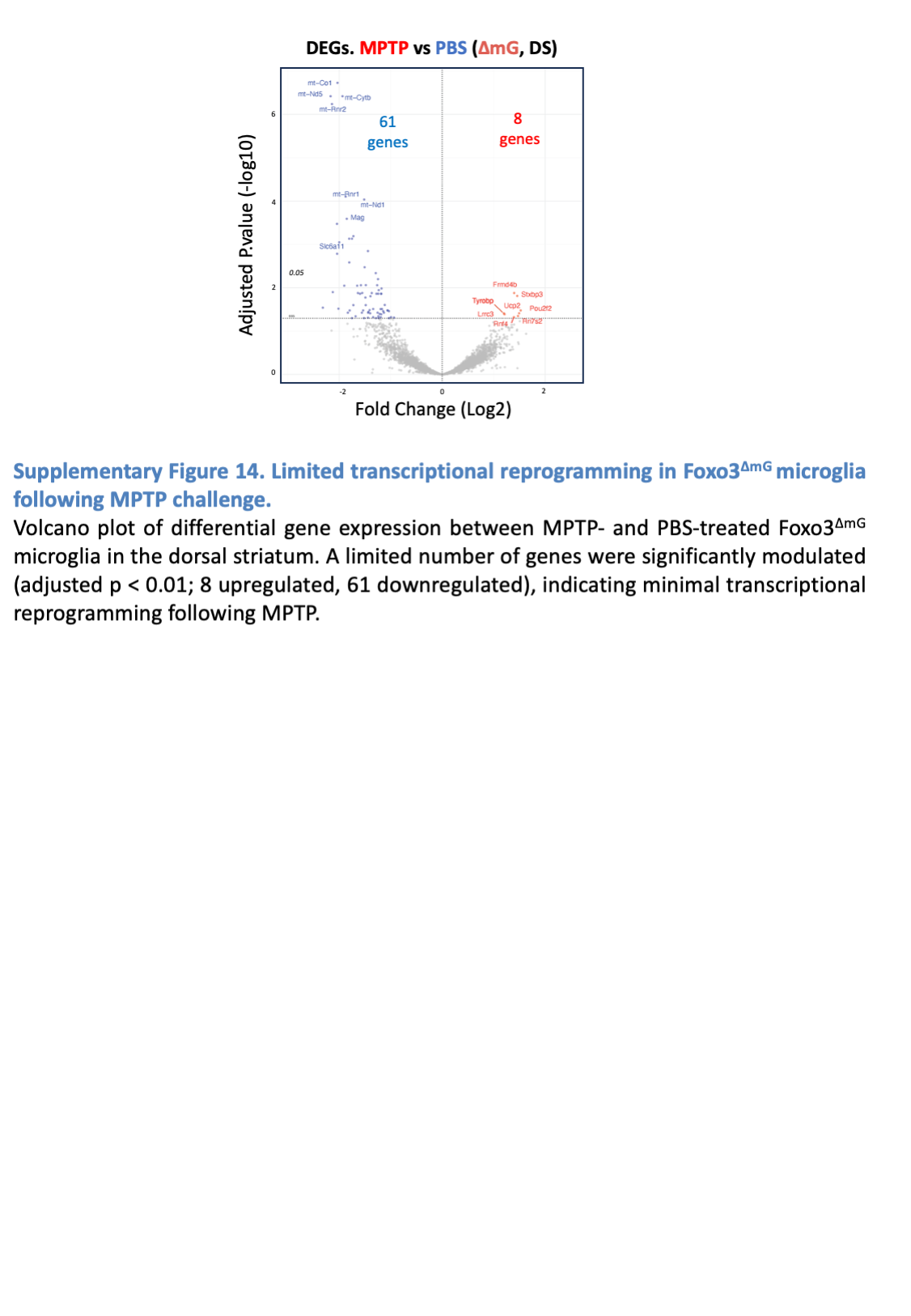

### SupFig15.png

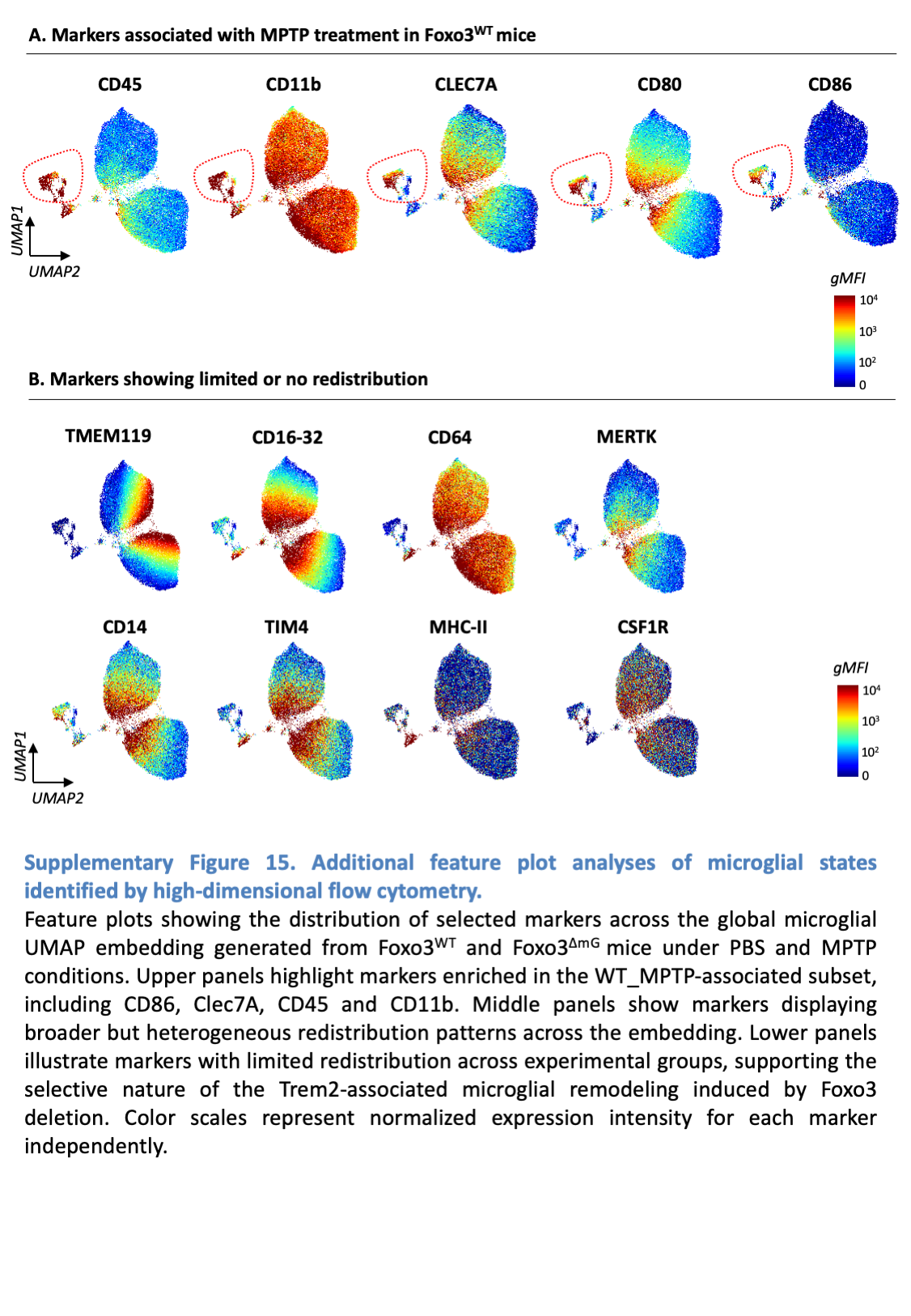

### SupFig16.png

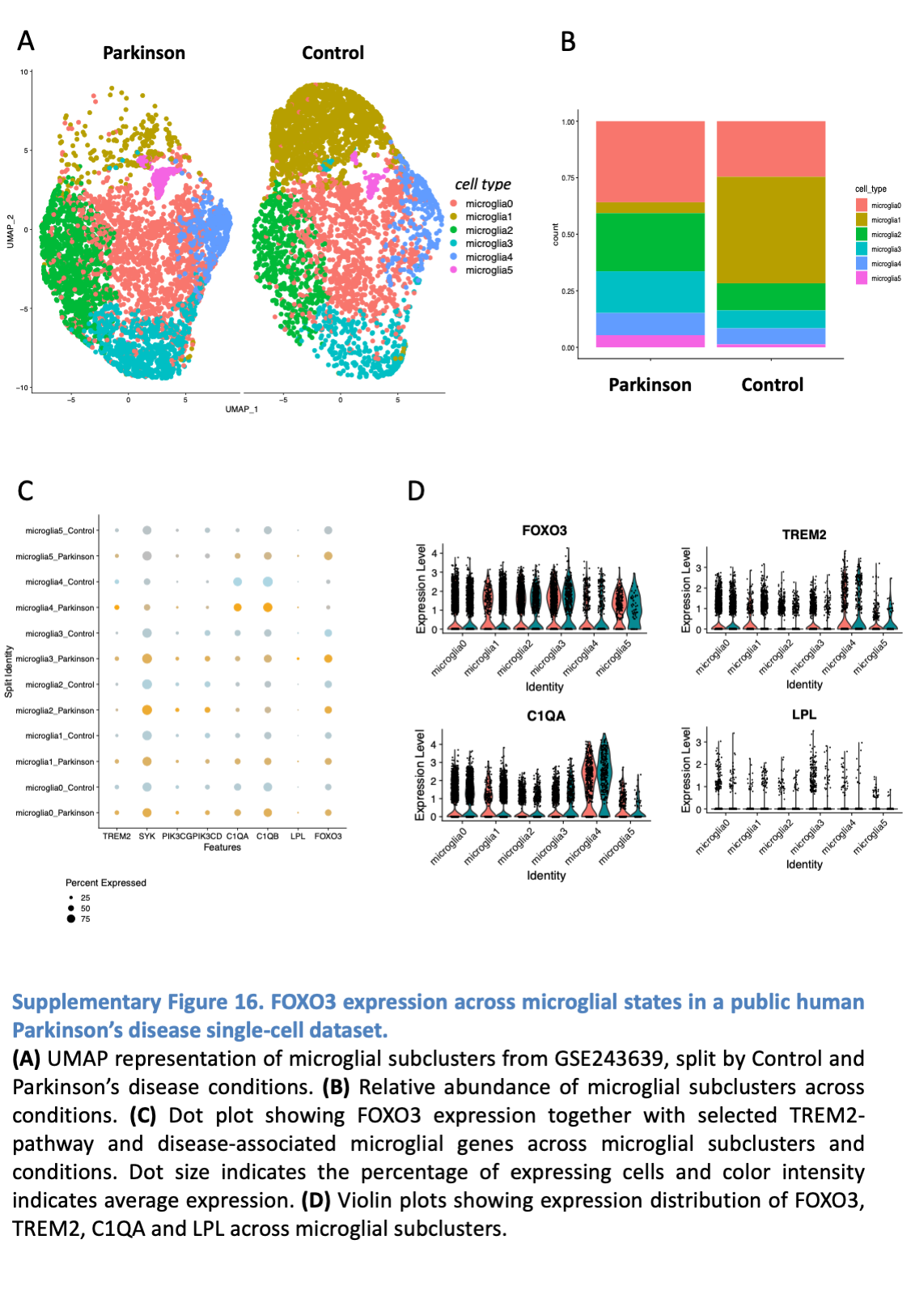
